# Genomic expansion of archaeal lineages resolved from deep Costa Rica sediments

**DOI:** 10.1101/763623

**Authors:** Ibrahim F. Farag, Jennifer F. Biddle, Rui Zhao, Amanda J. Martino, Christopher H. House, Rosa I. León-Zayas

**Author notes:** **Materials and Correspondence**, Correspondence and material requests should be addressed to Jennifer Biddle, < >.

## Abstract

Numerous archaeal lineages are known to inhabit marine subsurface sediments, although their distributions, metabolic capacities and interspecies interactions are still not well understood. Abundant and diverse archaea were recently reported in Costa Rica (CR) margin subseafloor sediments recovered during IODP Expedition 334. Here, we recover metagenome-assembled genomes (MAGs) of archaea from the CR-margin and compare them to their relatives from shallower settings. We describe 31 MAGs of 6 different archaeal lineages (*Lokiarchaeota, Thorarchaeota, Heimdallarchaeota, Bathyarcheota, Thermoplasmatales* and *Hadesarchaea*) and thoroughly analyze representative MAGs from the phyla *Lokiarchaeota* and *Bathyarchaeota*. Our analysis suggests the potential capabilities of *Lokiarchaeota* members to anaerobically degrade aliphatic and aromatic hydrocarbons. We show it is genetically possible and energetically feasible for *Lokiarchaeota* to degrade benzoate if they associate with organisms using nitrate, nitrite and sulfite as electron acceptors, which suggests a possibility of syntrophic relationships between *Lokiarchaeota* and nitrite and sulfite reducers. The novel *Bathyarchaeota* lineage possesses an incomplete methanogenesis pathway lacking the methyl co-enzyme M reductase complex and encodes a non-canonical acetogenic pathway potentially coupling methylotrophy to acetogenesis via the methyl branch of Wood-Ljundahl pathway. These novel metabolic characteristics suggest the potential of this *Bathyarchaeota* lineage to be a transition between methanogenic and acetogenic *Bathyarchaeota* lineages. This work substantially expands our knowledge about the metabolic function repertoire of marine benthic archaea.

## Introduction

Marine subsurface sediments are full of diverse archaeal lineages[1], although their distributions, ecological roles and adaptation strategies are still not well understood[2][3][4][5]. Metagenomic sequencing and single cell genomics have enabled the discovery of a great number of organisms, the elucidation of new metabolisms, the expansion of known lineages and the redefinition of portions of the tree of life[6][7], [8][9][10][11][12][13][14]. However, there is still a paucity of genomes resolved from the deep marine subsurface, meaning the niche specific adaptations in deep biosphere is not yet well understood.

Recently, the tree of life has been greatly expanded with the discovery of the *Asgard* superphylum, a deeply-branching monophyletic group thought to be some of the closest relatives to the eukaryotic branch of life[11][15]. Genome analyses of *Asgard* archaea have suggested diverse metabolic functions extending from an autotrophic lifestyle, primarily dependent on carbon fixation via Wood-Ljundahl pathway and acetogenesis, to a heterorganotrophic lifestyle consuming proteins and aliphatic hydrocarbons, using methyl-CoM reductase-like enzymes, to recycle aliphatic hydrocarbons released from the subsurface[16][17][18]. *Asgard* members were proposed to be engaged in symbiotic partnerships involving syntrophic transfers of hydrogen and electrons following the ‘reverse flow model’ [16].

Members of another archaeal phylum in the deep subsurface, *Bathyarchaeota*, are characterized by their wide metabolic repertoire enabling heterotrophic scavenging of proteins, carbohydrates, short chain lipids and other reduced compounds as substrates as well as their methane-metabolizing potential [19]. However, bathyarchaeotal genomes have also suggested the potential for carbon fixation and acetogenesis [20]. The evolutionary path describing the acquisition of both methanogenesis and acetogenesis pathways in *Bathyarchaeota* remains unresolved [20][21].

Deep sediment from the Costa Rica (CR) margin subseafloor, sampled during the International Ocean Discovery Program (IODP) Expedition 334, was recently shown to host abundant archaea [22]. Here we examine in detail the archaeal genomes recovered from the Costa Rica Margin and compare them to their relatives recovered from shallower sites. In this study, we report 31 archaeal metagenome-assembled genomes (MAGs) belonging to six different archaeal lineages (*Lokiarchaeota, Thorarchaeota, Heimdallarchaeota, Bathyarcheota, Thermoplasmatales* and *Hadesarchaea*) from the Costa Rica Margin subseafloor. We thoroughly analyze representative MAGs of two novel archaeal lineages belonging to phyla *Lokiarchaeota* and *Bathyarchaeota*. Our analysis suggests that *Lokiarchaeota* genomes encode for genes dedicated to process and degrade aliphatic and aromatic hydrocarbon anaerobically. We also describe a metabolically novel *Bathyarchaeota*, which lacks methyl co-enzyme M reductase (MCR) complex and possesses a non-canonical acetogenic pathway linking methylotrophy to acetogenesis via the methyl branch of Wood-Ljundahl pathway. Lastly, we integrate genomic and thermodynamic modeling to underline the ecological and physiological conditions that could drive the syntrophic interactions among CR-Asgards and the development of non-canonical acetogenesis in CR-*Bathyarchaeota*.

## Results

### Overall archaeal abundance and community structure

High abundances of archaea across sediment samples collected from five depths located along Costa Rica Margin (2, 32 and 93 mbsf from Site 1378 and 22 and 45 mbsf from Site 1379 of IODP Expedition 334), were previously reported[22]. Metagenomic data from these samples was assembled and examined for small subunit ribosomal genes. A total of 126 of 16S rRNA gene sequences were recovered from the metagenome assemblies. Sequences affiliated to archaea (31 sequences representing 25% of the total) reveal a wide diversity of lineages, including *Bathyarchaeota* (14 sequences representing 11 % of the total 16S rRNA gene sequences), *Thermoplasma* (5 sequences representing 4%) and *Lokiarchaeota* (4 sequences representing 3%) (Figure S1).

We compared two million raw metagenome reads (150 bp) for each dataset to the NCBI (nr) database [23]. Nearly 6-8% of the total short reads were successfully assigned to their respective phylogenetic group at the phylum level, while the phylogenetic signatures were not clear in the remaining reads (92-94%). The percentage of total prokaryote reads belonging to archaea ranged from 5% (at 93 mbsf, 1378) to 26% (at 32 mbsf, 1378).. Overall, community structure composition analysis conducted on metagenomic raw reads indicated the prevalence of two archaeal phyla in the Costa Rica margin: *Lokiarchaeota* and *Bathyarchaeota* (Figure 1B). *Lokiarchaeota* were most abundant among the archaea reads at the 32 mbsf samples in core 1378, while *Bathyarchaeota* reads were most abundant at 22 mbsf and 45 mbsf in core 1379.

**Figure 1.**
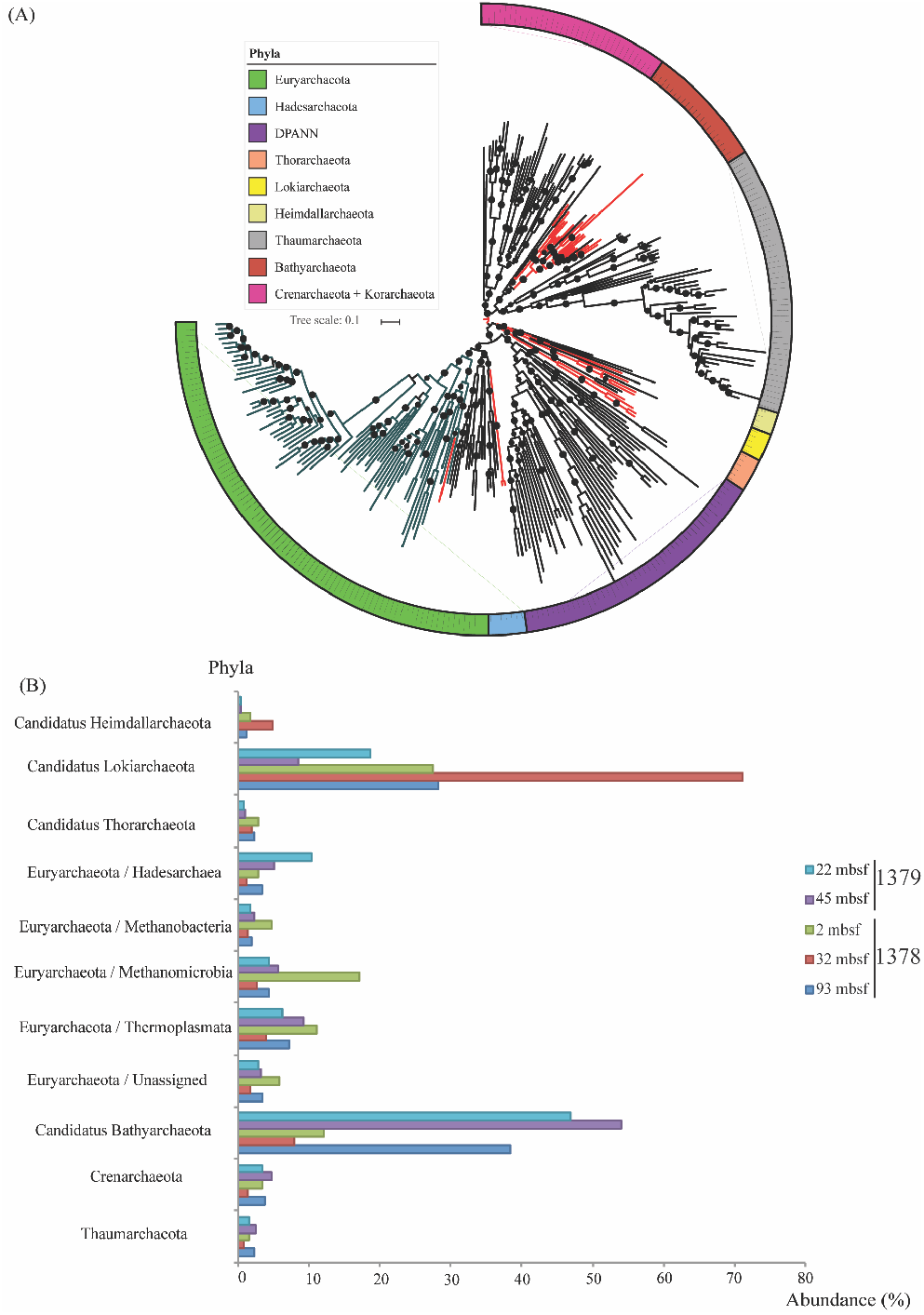
**(A) Phylogenetic placement of the Costa Rica archaeal draft genomes (genomes of this study are highlighted in red).** The maximum-likelihood phylogenetic tree was calculated based on the concatenation of 16 ribosomal proteins (L2, L3, L4, L5, L6, L14, L15, L16, L18, L22, L24, S3, S8, S10, S17, and S19) retrieved from the Costa Rica archaeal genomes and 231 reference archaeal genomes representing 13 different archaeal phyla. The relationships were inferred using the best fit substitution model (VT+F+R10) and nodes with bootstrap support >80% were marked by black circles. Scale bar indicates substitutions per site. The tree is available with full bootstrap values in Newick format in the Supplementary Data. **(B) Relative abundance percentages of all archaeal lineages making up > 1% of the total communities.** This graph was calculated by parsing the raw reads against NCBI (nr) database applying e-value cut off score 1e-5.

### General genomic features of the abundant archaeal lineages

Across the five-depths analyzed, 31 different draft archaeal MAGs were recovered (Table 1). Genomic analyses were only performed on the 11 MAGs showing high completeness and low contamination percentages (above 60% completeness and below 5% contamination). Overall, completeness varied from 32% to 99% with an average of 50% and contamination varied from 0 to 10% with an average of 8% (Table 1). Genome qualities were further assessed by comparing their predicted proteins against the NCBI (nr) database to evaluate the extent of phylogenetic consensus within the binned genomes (Figure 2A). Overall, the taxonomic affiliations of the majority of the predicted proteins (60-75%) in each genome agreed with their respective phylogenetic group, except CR-12 in which only 15% of the encoded proteins assigned to *Heimdallarchaeota* and was excluded from further analysis.

**Figure 2.**
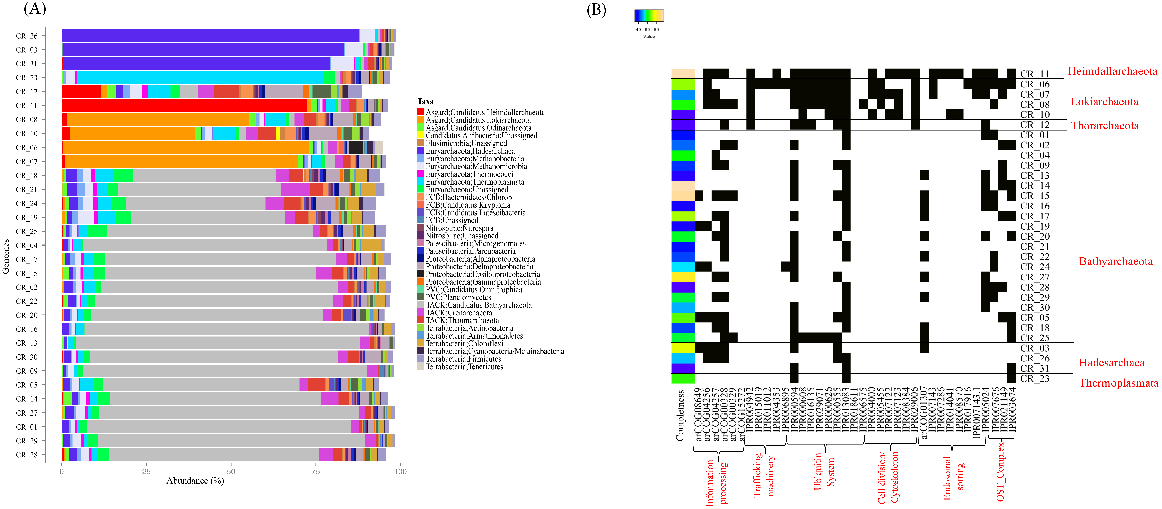
**(A) Phylogenetic distributions of the predicted proteins encoded by the archaeal MAGs, the phylogenetic assignments were performed through comparing the proteins to the NCBI (nr) proteins database via the DarkHorse software[23]. (B) Heat map shows the presence (black) /absence (white) patterns of eukaryotic homologs in the recovered archaea MAGs.** The color along the left side shows the completeness of the genome bin, as less complete bins would be expected to contain fewer homologs. Only the *Asgard* archaea contain cell division/cytoskeleton homologs.

**Table 1.** Details of the genomes constructed from the Costa Rica Margin

| Bin_ID | Dataset | Marker lineage | Phylogenetic affiliation | Completeness | completeness_metric | Contamination | Strain heterogeneity | Genome size (Mbp) | # scaffolds | GC | GC std (scaffolds > 1kbp) | Coverage |
| --- | --- | --- | --- | --- | --- | --- | --- | --- | --- | --- | --- | --- |
| CR_01 | 1378_2mbsf | k_Archaea (UID2) | Bathyarchaeota | 38.71 | partial | 10.09 | 59.09 | 0.62 | 335 | 39 | 6.02 | 155.6 |
| CR_02 | 1378_2mbsf | k_Archaea (UID2) | Bathyarchaeota | 45.64 | partial | 5.61 | 16.67 | 1.05 | 225 | 38.7 | 0 | 29.9 |
| CR_03 | 1378_2mbsf | p_Euryarchaeota (UID3) | Hadesarchaea | 74.86 | substantial | 10.67 | 50 | 0.98 | 333 | 40.2 | 6.68 | 19.3 |
| CR_04 | 1378_2mbsf | k_Archaea (UID2) | Bathyarchaeota | 60.62 | moderate | 10.08 | 5.88 | 1.09 | 368 | 42.7 | 4.79 | 16.05 |
| CR_05 | 1378_2mbsf | k_Archaea (UID2) | Bathyarchaeota (RPL2) | 65.2 | moderate | 10.26 | 0 | 1.92 | 900 | 40.9 | 0 | 11.4 |
| CR_06 | 1378_32mbsf | k_Archaea (UID2) | Asgard (Lokiarchaeota) | 86.78 | substantial | 1.87 | 33.33 | 2.41 | 1110 | 42.7 | 11.06 | 216.64 |
| CR_07 | 1378_32mbsf | k_Archaea (UID2) | Asgard (Lokiarchaeota) | 47.66 | partial | 5.43 | 63.64 | 1.525 | 829 | 41.9 | 0 | 72.54 |
| CR_08 | 1378_32mbsf | k_Archaea (UID2) | Asgard (Lokiarchaeota) | 61.06 | moderate | 9.65 | 20 | 3.54 | 693 | 42.9 | 8.76 | 71.06 |
| CR_09 | 1378_32mbsf | k_Archaea (UID2) | Bathyarchaeota | 41.64 | partial | 0 | 0 | 3.6 | 89 | 39.3 | 0 | 30.63 |
| CR_10 | 1378_32mbsf | k_Archaea (UID2) | Asgard (Lokiarchaeota) | 32.81 | partial | 10.6 | 23.33 | 2.91 | 1865 | 43.3 | 6.55 | 24.53 |
| CR_11 | 1378_32mbsf | k_Archaea (UID2) | Asgard (Heimdallarchaeota) | 94.39 | near | 9.06 | 0 | 3.47 | 167 | 38.6 | 7.88 | 23.11 |
| CR_12 | 1378_32mbsf | k_Archaea (UID2) | Asgard (Thorarchaeota) | 32.05 | partial | 1.08 | 0 | 1.49 | 1019 | 38.8 | 0 | 10.48 |
| CR_13 | 1379_22mbsf | k_Archaea (UID2) | Bathyarchaeota | 35.05 | partial | 0 | 0 | 0.3 | 118 | 39.2 | 0 | 116.39 |
| CR_14 | 1379_22mbsf | k_Archaea (UID2) | Bathyarchaeota | 95.64 | near | 4.15 | 0 | 2.49 | 140 | 39.4 | 7.77 | 77.12 |
| CR_15 | 1379_22mbsf | k_Archaea (UID2) | Bathyarchaeota | 92.61 | near | 3.12 | 0 | 2.11 | 130 | 39 | 5.31 | 47.44 |
| CR_16 | 1379_22mbsf | k_Archaea (UID2) | Bathyarchaeota | 36.14 | partial | 0 | 0 | 2.99 | 59 | 39.4 | 7.03 | 47.78 |
| CR_17 | 1379_22mbsf | k_Archaea (UID2) | Bathyarchaeota | 71.03 | substantial | 6.54 | 0 | 0.88 | 28 | 39.4 | 7.51 | 26.16 |
| CR_18 | 1379_22mbsf | k_Archaea (UID2) | Bathyarchaeota (RPL2) | 42.3 | partial | 5.61 | 0 | 1.13 | 103 | 40.8 | 6.46 | 24.29 |
| CR_19 | 1379_22mbsf | k_Archaea (UID2) | Bathyarchaeota | 35.84 | partial | 5.99 | 0 | 1.03 | 340 | 41 | 0 | 13.83 |
| CR_20 | 1379_22mbsf | k_Archaea (UID2) | Bathyarchaeota | 58.35 | moderate | 2.8 | 0 | 1.01 | 184 | 41 | 6.38 | 18.65 |
| CR_21 | 1379_22mbsf | k_Archaea (UID2) | Bathyarchaeota | 38.51 | partial | 1.94 | 0 | 0.58 | 249 | 41.8 | 5.06 | 17.31 |
| CR_22 | 1379_22mbsf | k_Archaea (UID2) | Bathyarchaeota | 42.88 | partial | 9.88 | 15.38 | 1.31 | 344 | 38.6 | 3.12 | 19.48 |
| CR 23 | 1379 22mbsf | p__Euryarchaeota (UID3) | Thermoplasmata | 62.44 | moderate | 2.4 | 0 | 1.96 | 284 | 43.1 | 5.55 | 19.88 |
| CR 24 | 1379 22mbsf | k__Archaea (UID2) | Bathyarchaeota | 53.74 | partial | 6.31 | 0 | 2.36 | 556 | 41.8 | 3.38 | 17.98 |
| CR 25 | 1379 22mbsf | k__Archaea (UID2) | Bathyarchaeota (RPS3) | 56.65 | moderate | 1.94 | 0 | 1.68 | 531 | 39.6 | 3.35 | 13.92 |
| CR 26 | 1379 22mbsf | k__Archaea (UID2) | Hadesarchaea | 50.96 | moderate | 7.79 | 0 | 0.58 | 329 | 39.6 | 0 | 14.9 |
| CR 27 | 1379 45mbsf | k__Archaea (UID2) | Bathyarchaeota | 79.77 | substantial | 5.49 | 33.33 | 1.24 | 404 | 39.8 | 4.19 | 63.85 |
| CR 28 | 1379 45mbsf | k__Archaea (UID2) | Bathyarchaeota | 32.78 | partial | 6.15 | 0 | 1.25 | 296 | 39.1 | 8.11 | 45.38 |
| CR 29 | 1379 45mbsf | k__Archaea (UID2) | Bathyarchaeota | 56.96 | moderate | 0.97 | 100 | 0.7 | 154 | 37.7 | 7.52 | 40.92 |
| CR 30 | 1379 45mbsf | k__Archaea (UID2) | Bathyarchaeota | 49.54 | partial | 3.4 | 28.57 | 4 | 191 | 40 | 0 | 21.23 |
| CR 31 | 1379 45mbsf | p__Euryarchaeota (UID4) | Hadesarchaea | 31.55 | partial | 0.64 | 33.33 | 3.75 | 176 | 40.2 | 0 | 25.74 |

Phylogenetic placement of the draft genomes was determined using 16 ribosomal proteins (Figure 1A and Figure S2)[12]. The recovered MAGs were affiliated to six different phylogenetic lineages, namely, *Lokiarchaeota*, *Thorarchaeota*, *Heimdallarchaeota*, *Bathyarcheota*, *Thermoplasmatales* and *Hadesarchaea*.

### Distribution of Eukaryotic signature protein (ESP) homologs in Costa Rica archaea MAGs

All archaeal MAGs recovered from Costa Rica sediments encoded eukaryotic signature proteins (ESPs). These ESPs include protein homologs dedicated for information processing, trafficking machineries, ubiquitin system, cell division and cytoskeleton formation. The *Asgard* archaea genomes recovered from CR, e.g. *Heimdallarchaeota* and *Lokiarchaeota* MAGs (CR_06, CR_11), showed significantly higher numbers of eukaryotic homologs and covered broader classes of ESPs (Figure 2b). However, ESPs were also detected in the *Bathyarchaeota, Hadesarchaeota* and *Thermoplasmata* MAGs recovered from CR (Figure 2b). Yet, only the *Asgard* genomes have the homologs for cell division and cytoskeleton.

### Genomic evidence of CR_Asgard hydrocarbon utilization

We screened all of the MAGS, assembled metagenomes and raw reads for the presence of genes encoding for MCR complex to assess the role of the archaeal members in mediating the degradation of hydrocarbon compounds (Figure 3A) that are abundant in Costa Rica (CR) margin [24]. MCR complex genes (*mcrABCDG*) were completely absent from both the MAGs and the entire metagenomes, which indicates that short chain alkanes are not oxidized using MCR complex in CR sediments. All metagenomic reads and MAGs were screened for possible alternative hydrocarbon degradation pathways using custom HMM searches specifically targeting key metabolic genes for aliphatic and aromatic hydrocarbons degradation pathways (Figure 3A). Multiple pathways were successfully identified including glycyl-radical enzymes (GREs) related genes coupled with n-alkane succinate synthase (AssA) and benzylsuccinate synthase (BssA), which activates *n*-alkanes and mono-aromatic compounds, respectively, by forming C-C bond between these compounds and fumarate to form hydrocarbon adducts [25] [26]. Benzylsuccinate synthesis is the initial step for aromatic hydrocarbon mineralization, in which benzylsuccinate is converted to benzoyl-CoA[25]. Interestingly, the capability of ATP-dependent Benzoyl-CoA reductase (BCR) complex utilization was identified in some CR *Asgard* members (*Lokiarchaeota, Thorarchaeota* and *Heimdallarachaeota*) (Figure 3). This reaction can dearomatize the benzoyl-CoA to dienoyl-CoAs as the first step in aromatic hydrocarbon degradation and then couple this reaction with a beta-oxidation pathway to ultimately produce acetyl-CoA, similar to the mechanism previously reported in the denitrifying bacteria *Thauera aromatica* [27]. Since *Lokiarchaeota* CR_06 had the least contamination levels <2%, we used this MAG to verify the presence of aromatic hydrocarbon degradation function mediated by BCR complex in *Asgards*. Phylogenetic analysis of the BCR subunit B recovered from *Lokiarchaeota CR_06* showed their affiliation to class Bzd, which is composed of four subunits (BzdONPQ) (Figure 3C). This BCR type was originally discovered in *Betaproteobacteria, Azoarcus evansii* and the ones detected in CR_06 is closely related to BCRs detected in different *β, δ proteobacteria*, and other archaeal lineages (e.g. *Lokiarchaeota, Bathyarchaeota* and *Archaeoglobus*) (Figure 3C)[28]. Subunits P and Q have ATP binding/ATPase functional domains and are linked together by a [4Fe-4S] cluster. Reduced ferrodoxins transfer electrons to the CoA-ester-binding domains (subunits O and N), which catalyzes the cleavage of benzoyl-CoA aromatic ring and yields dienoyl CoA product (Figure 3A) [28]. In CR_06, the four subunits of BCR complex were located in one contiguous operon (CR_06-contig-100_3495), however due to the high fragmentation of the genomes no reliable phylomarker genes were found in the adjoining genomic neighborhoods. A plausible explanation for the high fragmentation levels of the CR_06 (number of scaffolds= 1110), even though the genome exhibited high coverage levels >200x, is that there are high levels of intra-lineage strain heterogeneity. This is confirmed by the ANI values (ANI=70-80%) between MAGs of the same phylogenetic group and disparities in their coverage levels (table 1 and table S3).

**Figure 3.**
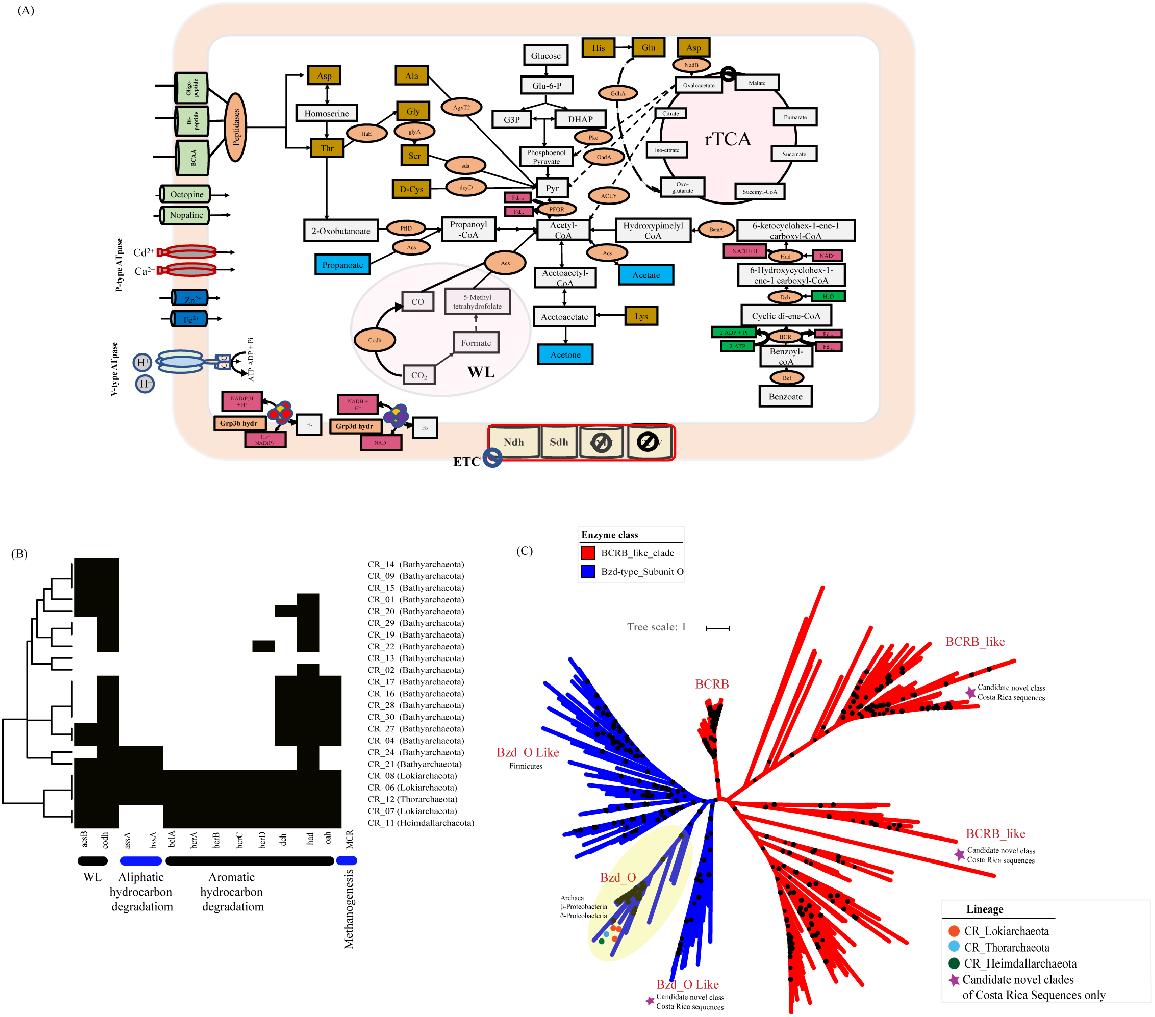
**(A) Metabolic reconstruction of the CR_Lokiarchaeota bin (CR_06).** Central metabolic pathways are shown in gray boxes, carbon fixation pathways (WL and rTCA cycles) are shown in pink, electron transport chain (ETC) proteins are shown in yellow, fermentation products are shown in blue boxes, amino acids are shown in brown boxes, enzymes and enzyme complexes are shown in orange circles, energy carriers are shown in red, energy molecules are shown in dark green, metabolite and amino acid transporters are shown in light green, and cation transporters are shown in dark blue. **(B) Distribution patterns of the hydrocarbon degradation pathways among the constructed CR-Archaea genomes.** Genomes were clustered based on presence (black) /absence (white) profiles using Euclidean distance and average linkage method. X axis represents enzymes included in the analysis, bottom bars are the pathways that these enzymes represent and y axis includes bin names/phylogenetic affiliations. **(C) Maximum likelihood tree of the benzoyl-CoA reductase subunit B.** The tree was calculated using the best fit substitution model (VT+F+R7) that describes the evolutionary relationships between BCRB families. The tree was made using reference sequences under the KEGG entry (K04113) collected from AnnoTree[30] and branch location was tested using 1000 ultrafast bootstraps and approximate Bayesian computation, branches with bootstrap support >80% were marked by black circles. Blue and Red clades highlight sequences belong to Bzd_O and BCR_B subfamilies, respectively. Scale bar indicates substitutions per site. Sequences from CR*_Lokiarchaeota* bins was marked with red circles, *CR_Thorarchaeota* bin was marked with light blue and *CR_Heimdallarchaeota* was marked with green. Candidate novel clades present in the Costa Rica metagenomic datasets were marked with purple stars. The tree is available with full bootstrap values in Newick format in the Supplementary Data.

The affiliations of the BCR complex related proteins, BCRA and BCRB in *Lokiarchaeota* CR_06 were confirmed by the following observations: 1) contigs encoding for the four subunits of BCR complex fall within the same coverage and GC% range of the rest of the contigs in the same MAG (Figure S3); 2) phylogenetic trees of BzdO and Q protein sequences detected in the CR_06 were placed as siblings to BCR sequences belonging to *Lokiarchaeota* MAGs recovered from deep sediment from south China Sea (GenBank accession numbers TET61664 and TKJ21612) (Figure 3C, S4)[29].

### Fate of the degraded aromatic hydrocarbons and syntrophic interactions

CR_ *Lokiarchaeota (CR_06*) as well as other *Asgard* in the CR margin could potentially grow heterotrophically on aromatic hydrocarbons. Though, the absence of genes encoding for different types of cytochrome oxidase and anaerobic respiration from the genome content of the *Asgard* MAGs (CR_06, 07, 08, 11 and 12) indicate their inability to completely mineralize these hydrocarbons to CO_2_ and H_2_O, which would allow a high energy yield. They encode for the genes mediating the fermentation of these organic macromolecules to acetate and other reduced products, which is thermodynamically unfavorable under CR conditions with a positive ΔrG’ value (ΔrG’= 196.3 [kJ/mol]) (Figure 3 and table S1), suggesting that these genes are maintained due to a thermodynamically favorable force. Due to the genome incompleteness (86% complete), we cannot rule out the possibility for the presence of complete aromatic hydrocarbon mineralization pathways using one or more of oxidized substrates as electron sinks (Table 1). More likely, however, *Lokiarchaetoa* are gaining energy through syntrophic interactions with partners capable of oxidizing the biodegradation intermediates.

Here, we inferred the identity of the potential syntrophic partners under marine subsurface conditions by comparing all the possible metabolic and thermodynamic scenarios, gauging each scenario based on the presence of the metabolic pathways in our metagenomic datasets and the thermodynamic feasibility under each condition (Figure 3B, Table S1 and S6). We calculated the Gibbs free energy of coupled reactions under a wide range of substrate concentration conditions: Reaction 1-5 (Table S1, Figure S5). Best conditions suggest that the degradation of benzoate (the central metabolic intermediate in aromatic hydrocarbon degradation pathways) potentially occurs under the following metabolic conditions (a, b, and c): (a) benzoate mineralization to CO_2_ and H_2_O coupled with nitrite reduction to ammonia (ΔrG’= −1206.3 [kJ/mol]); (b) benzoate mineralization to CO_2_ and H_2_O coupled with sulfite reduction to hydrogen sulfide (ΔrG’= −373.6 [kJ/mol]); and (c) benzoate mineralization to CO_2_ and H_2_O coupled with nitrate reduction to nitrite (ΔrG’= −119.9 [kJ/mol]).

The type and complexity of the exchanged substrates are another key factor that may shape the syntrophic relationship and the identity of the syntrophic partners of *Lokiarchaeota*.. The presence of genes encoding for membrane bound electron bifurcating classes of [NiFe] hydrogenases, groups 3b and 3d, which couple the oxidation of NADH+ and NADPH+ with H_2_ evolution (Figure S6)[31] also suggests there is syntrophic exchange of hydrogen between *Asgard* and their partners. Additionally, we located genes encoding for β-oxidation enzymes (enoyl-CoA hydratase, acyl CoA dehydrogenase, and acetyl CoA acetyl transferase) and various fermentation pathways (acetate and formate)) in *Lokiarchaeota* CR_06, which suggests that short chain fatty acid and different fermentation products could also be syntrophically exchanged (Table S4). The diverse nature of substrates could facilitate the interactions between a broader range of partners of diverse metabolic capabilities and support the conclusions driven from our thermodynamic calculations.

It is worth noting that efficient substrate and electron exchange between syntrophic partners require the presence of either biological conduits (e.g. type IV pili or flagella) or some sort of electron shuttles allowing extracellular electron transfers (e.g. multiheme cytochromes) [32][33]. Hence, we screened CR_06 for these mechanisms and identified two candidate mechanisms for interspecies substrate and electron exchange. First, flagellar proteins are encoded by the CR_06, suggesting flagella as a potential structure mediating inter-species interactions. Second, CR_06 harbors a gene encoding for an oxidoreductase belonging to electron transfer flavoprotein-quinone oxidoreductase (CR_06_contig-100_4953_2), ETF-QO/FixC family, which potentially mediating the transfer of electrons across membranes.

### Other metabolic features of Lokiarchaeota (CR_06)

The genomic analysis of *Lokiarchaeota* (CR_06) MAG also suggests versatile catabolic capacities potentially targeting detrital proteins and short chain fatty acids (e.g. propanoate, butyrate), which are abundant in benthic marine sediments. CR_06 MAG has a relatively large number of peptidases encoding genes (92 peptidases/1Mbp) with diverse catalytic residues (e.g. aspartic, metallo, serine, etc.), which potentially degrade detrital proteins (Figure S7). It also contains genes encoding for various classes of facilitated and active transporters, which are dedicated to shuttle oligo/di-peptides and single amino acids (e.g. polar and branched chain amino acids) across the cell membrane. Also, CR_06 encoded for enzymes enabling the utilization of wide range of amino acids (e.g. aspartate, threonine, alanine, glycine, serine, cysteine, histidine, glutamine, and lysine), channel them to the central metabolic pathways and ultimately produce energy via fermentation (Figure 3A).

CR_06 MAG suggested the capacity to break down short chain fatty acids e.g. propanoate, oxo-butanoate as other potential substrates. Both propanoate and oxo-butanoate are converted to propanoyl CoA via formate acetyl transferase and acetyl synthase, respectively. Then, the resulting propanoyl CoA is converted to acetyl-CoA via malonyl-CoA pathway. CR_06 as well as other *Asgards* showed different autotrophic capacities enabling carbon fixation to complex organic carbon compounds using both the Wood-Ljundahl (WL) pathway and the reverse tricaboxylic acid (rTCA) cycle (Figure 3A).

### Metabolic features of CR-Bathyarchaeota

The other main lineage of Archaea in these sediments is *Bathyarchaeota. Bathyarchaeota* CR_14 is described in details since it has the highest quality at >95% complete and 4% contaminated. Phylogenomic analysis of *Bathyarchaeota* MAGS showed that CR_14 is clustered together with other CR*_Bathyarcaheota* in two distinct clades within *Bathyarchaeota* phylum (Figure S2). Also, relatively low similarity scores were observed between CR_14 and other reference *Bathyarchaeota* MAGs, which suggest that CR_14 belongs to a novel *Bathyarchaeota* class (Table S2).

Metabolic analysis revealed that CR_14 harbors genes encoding for incomplete methylotrophic methanogenesis pathway, yet is missing the key genes encoding for MCR complex (*mcrABCDG)*. The absence of this complex suggests that CR_14 is incapable of methanogenesis and methane metabolic genes may be rewired to perform different functions, where they recycle methyl groups from different methylated compounds and replace the functions of methyl branch in Wood-Ljungdahl pathway (Figure 4-A). CR_14 showed the potential capability to use formate as electron and hydrogen donor through using Group 4 hydrogenases (formate hydrogenlyases) (Figure S6)[31], and these electrons reduce trimethylamine N-oxide (TMAO) to trimethylamine via trimethylamine-oxide reductase or anaerobic dimethyl sulfoxide reductase (TMAO/DMSO reductase)[34]. Together the presence of genes encoding for trimethylamine-specific corrinoid protein as well as diverse classes of methyltransferases, CoB—CoM heterodisulfide reductase/ F420 nonreducing hydrogenase (hdrABCD and mvhADG) suggest the capability of CR_14 to recycle coenzyme M (CoM) and coenzyme B (CoB) and transfer the methyl group from trimethylamine to CoM-SH[20][35]. We located genes encoding the tetrahydromethanopterin S-methyltransferase (mtrA-H), suggesting that mtrA protein transfers the methyl group from CoM to 5,6,7,8-Tetrahydromethanopterin (H4MPT) and assimilates the methyl group into acetyl-CoA via the beta subunit of CODH/ACS complex, replacing the function of the methyl branch of Wood-Ljungdahl pathway. This agrees with our finding that only the genes encoding for the carbonyl branch of Wood-Ljundahl coupled with acetate fermentation genes (acetogenesis) were present in CR_14 and the genes encoding for methyl branch were completely missing for the same pathway. These collective metabolic features in *Bathyarchaeota* CR_14 suggest this genome may be a *Bathyarchaeota* lineage that bridges the gap between methanogenic and acetogenic *Bathyarchaeota* through adopting a non-canonical acetogenic life style (Table S5) [20][21].

**Figure 4.**
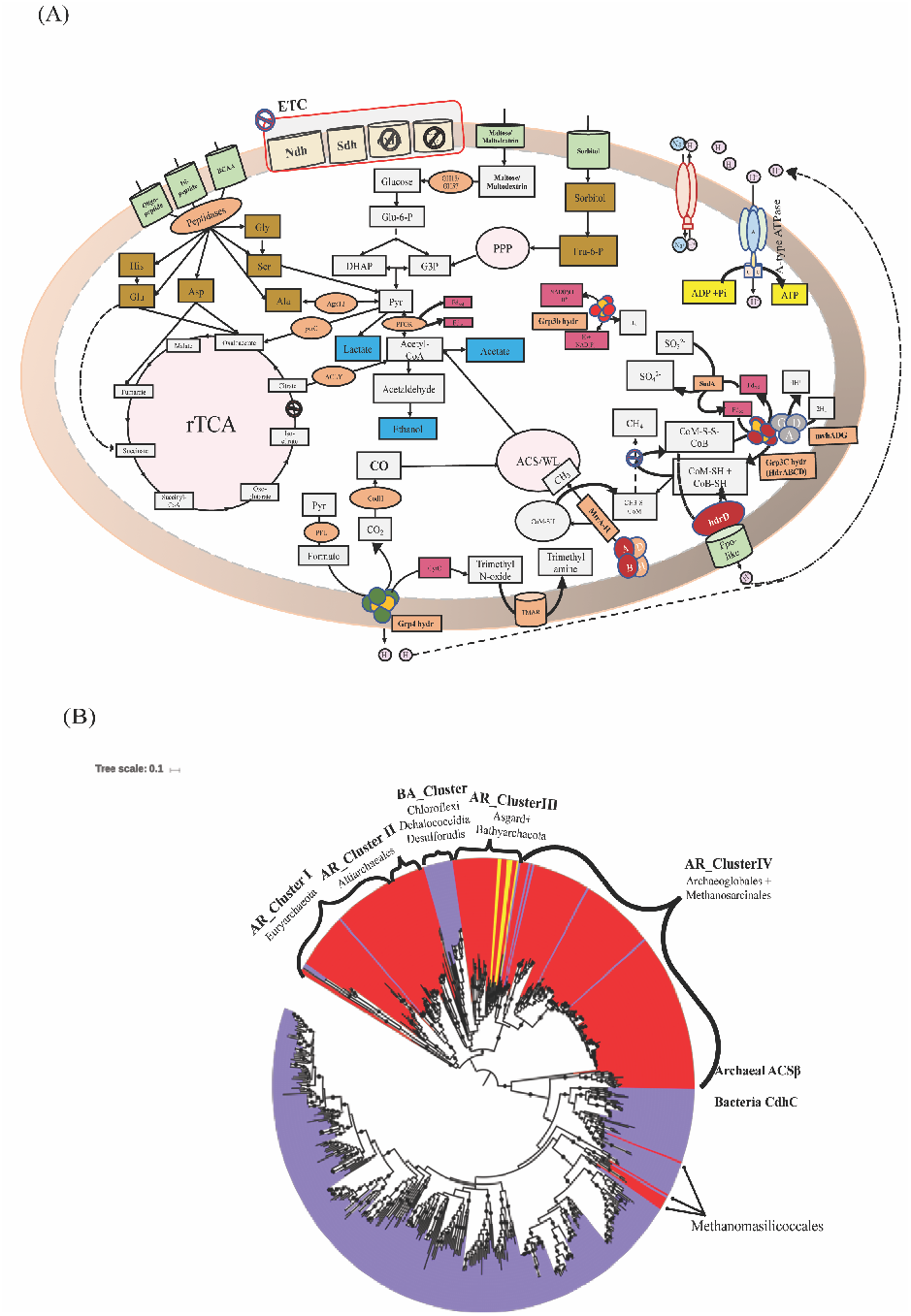
**(A) Metabolic reconstruction of the Bathyarchaeota bin CR_14.** Central metabolic pathways found in the genome (glycolysis, carbonyl branch of Wood-Ljundahl, and methanogenesis related genes) are shown in gray boxes, carbon fixation pathways (ACS/WL, PPP and rTCA cycles) are shown in pink, electron transport chain (ETC) proteins are shown in yellow, fermentation products are shown in blue boxes, amino acids are shown in brown boxes, enzymes and enzyme complexes are shown in orange circles, energy carriers are shown in red, metabolite and amino acid transporters are shown in light green. **(B) Maximum likelihood tree of the acetyl CoA synthase β subunit (ACSβ/CdhC).** The tree was calculated using the best fit substitution model (LG + R9) that describes the evolutionary relationships between ACS families. The tree was made using reference sequences under the KEGG entry (K00193) collected from AnnoTree[30] and branch location was tested using 1000 ultrafast bootstraps and approximate Bayesian computation, branches with bootstrap support >80% were marked by black circles. Blue and Red clades highlight sequences belong to bacterial (CdhC) and archaeal (ACSβ) versions, respectively. Scale bar indicates substitutions per site. Sequences from *CR_Lokiarchaeota* and *CR_Bathyarchaeota* bins were shaded with yellow. The tree is available with full bootstrap values in Newick format in the Supplementary Data.

Although the proposed scenario where CR_14 performs acetogenesis instead of methanogenesis is metabolically feasible, it is not clear why CR_14 invests a large amount of energy to maintain genes for methane metabolism. A plausible explanation for expressing methanogenesis-related genes is the lack of dedicated methyltransferases that could transport methyl groups directly between methylated compounds (e.g. trimethylamine) to Tetrahydromethanopterin (H4MPT). Therefore, encoding genes mediating the synthesis and cycling of CoM is necessary to use it as intermediate carrier to transport methyl groups from/to H4MPT. Also, the phylogenetic tree of the acetyl-CoA synthase beta subunit (AcsB) showed that CR_14 AcsB genes clustered together with genes recovered from other *Bathyarchaeota* lineages, *Asgards, Chloroflexi* and *Altiarchaeales* (Fig 4-B). The uniqueness of that clade stemmed from the previous hypothesis that *Altiarchaeales* members possess an ancient version of AcsB [36]. Likewise, members of *Chloroflexi* encoded for this archaeal version of ACSβ subunit similar to the ones present in *Bathyarchaeota, Asgards*, and *Altiarchaeales* [37][38]. Finally, this ACS type supports the hydrogen dependent, autotrophic life style of *Asgards* [16]. Accordingly, we hypothesize that *Bathyarchaeota* possess the type of ACS capable of incorporating methyl groups into acetyl-CoA utilizing different carrier proteins (CoM, H4MPT and may be other unknown carriers), allowing *Bathyarchaeota* to assimilate methyl groups originated from various methylated compounds.

## Discussion

In this study, we employed a metagenomics-enabled genomics approach to recover representative MAGs from the abundant archaeal phyla inhabiting deep sediments of Costa Rica margin and elucidate their potential ecological roles. A total of 31 MAGs belonging to archaeal phyla (*Lokiarchaeota, Thorarchaeota, Heimdallarchaeota, Bathyarcheota, Thermoplasmatales* and *Hadesarchaea*) were successfully recovered from five metagenomic datasets representing five different samples. Only 11 MAGs met our completion and contamination thresholds, >60% complete and <10% contamination, to be considered for the detailed genomic analyses and metabolic reconstruction. More than 90% of the high-quality genomes were affiliated to *Bathyarchaeota* and *Asgard* and the phylogenetic affiliations of the predicted proteins in each MAG confirmed the proper quality of the MAGs considered in this study. Remarkably, all the CR archaeal MAGs were enriched with ESP encoding genes. The wide distribution of these eukaryotic homologs indicates that ESPs are more ubiquitous in anaerobic archaea than previously recognized.

Notably, the sediments used in this study were collected from much deeper sites compared to the sediment where previously reported *Bathyarchaeota* and *Lokiarchaeota* were found. As such, our analysis was focused on MAGs belonging to *Bathyarchaeota* and *Lokiarchaeota* to try to understand their ecological potentials under these deep marine sediment conditions.

### Community interactions and Lokiarchaeota metabolic interdependencies

Deep marine sediments are rich in aliphatic and aromatic hydrocarbons that provide the associated microbes with a significant portion of their energy and carbon needs [39][40]. Previous studies showed that hydrocarbon degradation is restricted to limited bacterial and archaeal phyla (e.g. *Aminicenantes, TA06, Aerophobetes, Atribacteria, Helarchaeota* and *Bathyarchaeota*) [18][20] [41]. However, the full spectrum of archaea involved in the hydrocarbon degradation processes and the nature of their interactions are not fully elucidated. In the light of the above evidence, this study provides new views regarding the ecological roles and potential metabolic capacities of *Lokiarcaheota*. CR*_Lokiarchaeota* genome analysis expanded the range of metabolic features encoded by this phylum and predicts metabolic functions enabling the utilization of aliphatic and aromatic hydrocarbons as carbon and energy sources. Considering the high energy demands for aromatic hydrocarbon breakdown under the energy limited conditions of subseafloor sediments [28], it was unexpected to find the ATP-dependent BCR complex (class Bzd) in CR-*Asgards*/CR*-Lokiarchaeota* genomes while they have fermentation and/or acetogenic life styles. This strongly suggests that acetogenesis or acetate fermentation cannot be the ultimate fate of the aromatic hydrocarbon degradation process due to the low energy yields of these pathways. Under these circumstances, we propose that the CR-*Lokiarchaeota* members have the capacity of completely mineralizing aromatic hydrocarbons in syntrophy with microbes capable of using nitrite, sulfite and nitrate as electron sink under the subsurface settings to increase their energy budgets and sustain their energy requirement. After testing all possible partnership scenarios based on the presence/absence profiles of the candidate pathways and the thermodynamic feasibility (Figure 5 A-B), we found many potential partners for *Lokiarchaeota* belong to diverse metabolic groups (e.g. nitrate reducers, nitrite reducers, sulfate reducers, sulfite reducers and thiosulfate reducers), however only sulfite, nitrate and nitrite reducers are thermodynamically favored. In the same context, a diverse group of syntrophs might require sharing substrates of different qualities (e.g. acetate, propanoate, and other short chain fatty acids). Accordingly, this suggests the presence of *Lokiarchaeota* syntrophic partners capable of acetate oxidation and short chain fatty acid oxidation in addition to the hydrogenotrophic ones as previously proposed [16]. Therefore, we hypothesize that thermodynamic favorability and potential diversity of the shared metabolites will push the *Lokiarchaeota* to syntrophy, beyond individual metabolite exchange.

**Figure 5.**
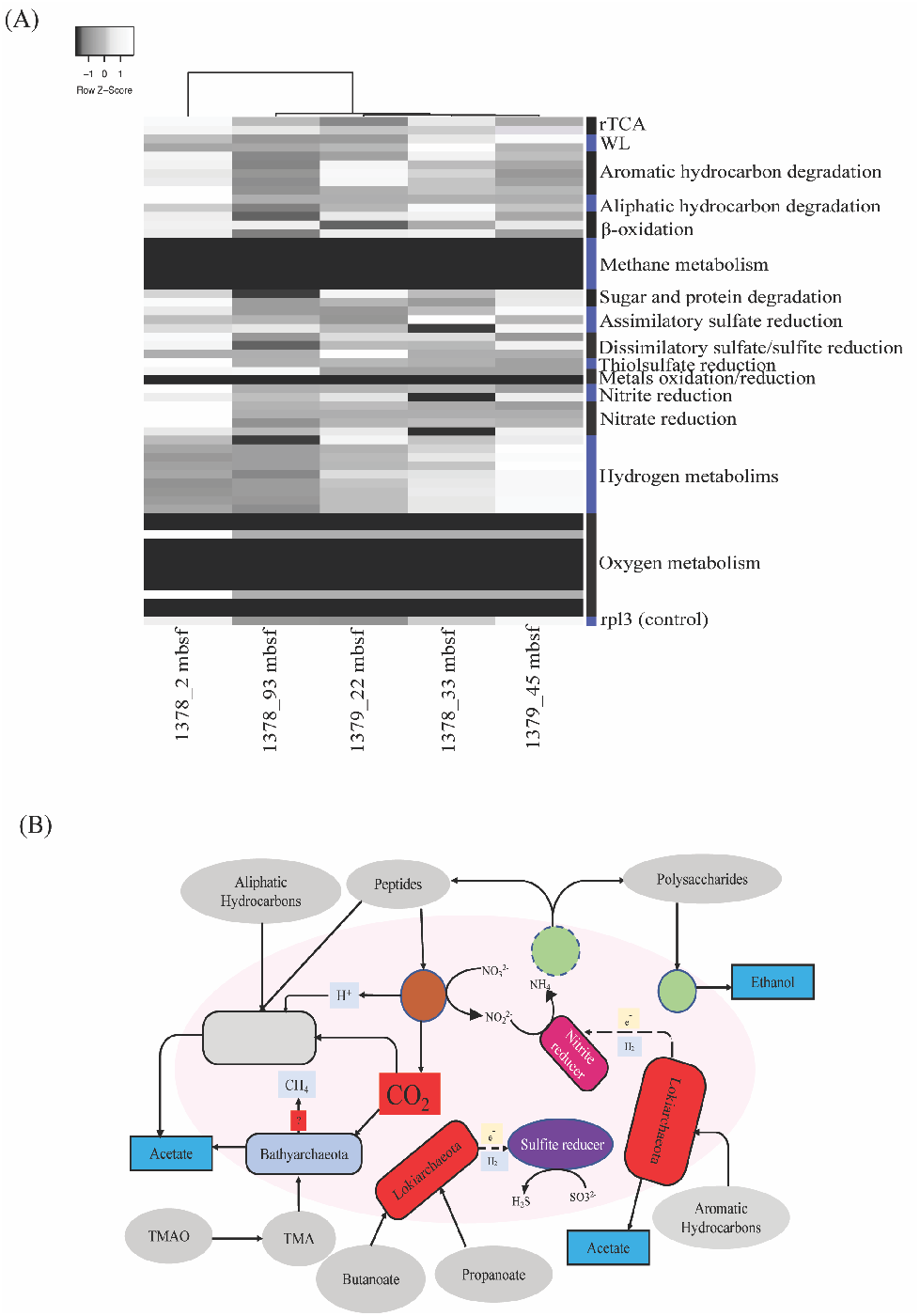
**(A) Heat map depicts the distribution patterns of the key metabolic pathways across all sampled metagenomic datasets.** Metagenomes were clustered based on presence (black) /absence (white) profiles using Euclidean distance and average linkage method. Y axis represents pathways included in the analysis and x axis includes metagenomic datasets analyzed. **(B) Hypothetical models for the microbial interactions in the Costa Rica sediments.** The potential substrates degraded by *Lokiarchaeota, Bathyarchaeota* and other microbes in the environment were colored in grey, potential metabolic products (in blue), potential interactions between *Lokiarchaeota* and their syntrophic partners and possible shared metabolites

### CR-Bathyarchaeota bridge the gap between acetogenic and methanogenic lineages

*Bathyarchaeota* have been shown to have both methanogenic and acetogenic lifestyles [20][21], yet the evolutionary trajectory and the ecological context driving the switch between lifestyles are not fully understood. Generally, *Bathyarchaeota* lineages are widely distributed in different benthic marine habitats compared to methanogenic ones [19], which could be explained by their acetogenesis capacity that enables the degradation of wide range of organic compounds under thermodynamic favorable conditions [42].

Interestingly, phylogenomic analysis revealed that *CR_Bathyarchaeota* MAGs are potentially affiliated to novel class that are significantly divergent from previously reported acetogenic and methanogenic lineages. In these CR sediments, distinct environmental conditions may have captured the intermediate transition between acetogenic and methanogenic lifestyles. We propose that these environmental conditions include the presence of high levels of methylated compounds (e.g. methylated amines) in deep-sea environments [43] and the absence of dedicated pathways/carriers necessary to recycle these methylated compounds and directly shuttle the extracted methyl groups to the Wood-Ljungdahl pathway. Also, there is a relatively high redox potential and substantial abundance of oxidized substrates that could serve as terminal electron acceptors in CR sediments, this may not favor the occurrence of full methanogenesis and the associated methanogenic archaea, as indicated by the absence of *mcrA* genes at the sampled depths (Figure 5A).

In summary, this study presents a large dataset of subsurface archaeal MAGs, some of which are high completeness and quality. Numerous archaeal MAGs host eukaryotic signatures, yet, only the *Asgard* genomes have the homologs for cell division and cytoskeleton. We also report a metabolically novel *Lokiarchaeota* lineage capable of aliphatic and aromatic hydrocarbon degradation with a putative partnership with metabolically diverse syntrophic organisms. Also, we revealed the presence of intermediate stage between acetogenic and methanogenic *Bathyarchaeota* that could convert methylated amines to acetate through linking methylotrophy to acetogenesis.

## Materials and Methods

### Site information and Sample collection

Samples used in this study were collected under aseptic conditions from Sites U1378 and U1379 of Costa Rica Margin during IODP Expedition 334. The sample depths are in the range of 2 – 93 meters below seafloor (mbsf). Detailed site descriptions were previously reported in the IODP Proceedings for Expedition 334 [44][22].

### DNA Extraction and Sequencing and genomes binning

DNA extraction and metagenomic sequencing have been described previously [22]and data have been deposited at NCBI GenBank SRA under project PRJEB11766. Metagenomic reads were quality trimmed using Nesoni following default parameter and applying q20 for quality score (www.vicbioinformatics.com/software.nesoni.shtml). Quality-controlled reads in individual samples were assembled separately using IDBA-UD[45] with default settings. Contigs longer than 1 kb were binned into MAGs using MaxBin V2.2.7[46]and further curated manually using VizBin[47]and through filtering outlier scaffolds not falling within the same GC% and differential coverage levels across different Costa Rica datasets. The quality and completeness of the MAGs were assessed using CheckM (v.1.0.7) [48]. MAGs from all give samples were dereplicated using dRep (version v2.0.5 with ANI > 99%)[49] and most complete MAG per taxon was selected for downstream analyses. Archaeal bins were further analyzed to determine their phylogenetic placements through the analysis of single copy marker genes using Phylosift[50] and 16 ribosomal proteins (see description below). Assembled contigs larger than 1kb were annotated using PROKKA[51]. Encoded proteins were predicted using Prodigal v2.6.3 with the default translation table (table 11) was applied[52].

### Concatenated ribosomal protein phylogeny

A maximum-likelihood tree was calculated based on the concatenation of 16 ribosomal proteins (L2, L3, L4, L5, L6, L14, L15, L16, L18, L22, L24, S3, S8, S10, S17, and S19) using IQ-Tree[53] (located on the CIPRES web server)[54]. References sequences used were collected [12] with the addition of more representatives of the *Bathyarchaeota* and *Asgard* lineages. Evolutionary distances were calculated based on best fit substitution model (VT+F+R10), and single branch location was tested using 1000 ultrafast bootstraps and approximate Bayesian computation[55][53], branches with bootstrap support >80% were marked by black circles.

### Metabolic reconstruction and Functional annotation

Predicted proteins from all MAGs were screened using HMMsearch tool against custom HMM databases representing the key genes for specific metabolic pathways [56]. The completion of the pathways was assessed through querying the predicted proteins against KEGG database using BlastKoala tool[57]. Carbohydrate-active enzymes (CAZymes) were identified using dbCAN-fam-HMMs (v6) database[58]. Cellular localizations of predicted proteins were identified using SignalP v5.0[59]. Proteases, peptidases, and peptidase inhibitors were identified using USEARCH-ublast tool[60] against the MEROPS database v12.1[61]. Transporters were identified using USEARCH-ublast tool[60] against the TCDB database[62]. Eukaryotic signature proteins were detected using Interpro v75.0[63]. Phylogenetic distributions of the predicted proteins in each bin were detected through comparing these proteins against the NCBI (nr) protein database via the DarkHorse software[23].

### Functional proteins-based trees

All functional proteins-based trees were built by aligning the query proteins sequences to the reference sequences belonging to the same protein family using Muscle v3.8.31[64]. Reference sequences were collected from AnnoTree[30] using the corresponding KEGG entry as search keyword. Aligned sequences were manually curated using Geneious v9.0.5 (https://www.geneious.com). The phylogenetic trees were computed using IQ-TREE (v1.6.6) [53], through the CIPRES web server[52] and the evolutionary relationships were described using the best fit model. Branch locations were tested using 1000 ultrafast bootstraps and approximate Bayesian computation[55][53].

### Thermodynamic calculations

Initial Gibbs free energy (ΔrG’) calculations were performed for the redox reactions proposed in the CR *Lokiarchaeota* and their poteintial partners using eQuilibrator[65] applying pH 8 similar to the approach reported in[42]and reactant concentrations 1mM each.

We further confirmed if the coupled redox reactions proposed for the novel CR *Lokiarchaeota* genomes were feasible, we calculated the Gibbs free energy of five redox reactions (Table S1) under the near in situ conditions at CR core U1378 and U1379, following the method described in LaRowe and Amend (2015)[66]. Gibbs free energy was calculated using the equation:

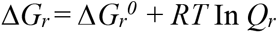

where 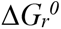 and *Q_r_* refer to the standard molar Gibbs energy and the reaction quotient of the indicated reaction, respectively, *R* represents the gas constant, and *T* denotes temperature in Kelvin. In this study, 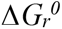 was calculated using the thermodynamic data of standard Gibbs free energy of formation of each species and corrected to near *in situ* pressure and temperature (4°C), using the *R* package *CHNOSZ* [67]. *Qr* stands for the reaction quotient, which can be calculated with the relation

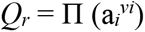

where a_*i*_ is the activity of species *i* and *v_i_* is its stoichiometric coefficient. a_*i*_ is the product of chemical species concentration [i] and its activity coefficient *γ_i_*, which was computed as a function of temperature and ionic strength by using an extended version of the Debye-Huckel equation[68]. Because most of the reactant concentrations were hard to measure or were below detection limits, we assumed 0.1 μM for the concentration of NO_3_^-^, NO_2_^-^, SO_3_^2-^, and H_2_S. We considered a wide range of concentrations for benzoate (C_7_H_5_O_2_^-^) (0.0001 – 100 μM), to explore the feasibility of these reactions over a wide range of substrate concentration changes.

## Acknowledgements

This work was funded by IODP 334 Post Expedition award T334A40, Exxon Mobil Research and Engineering, Penn State Astrobiology Research Center (through the NASA Astrobiology Institute, cooperative agreement #NNA09DA76A), the Pennsylvania Space Grant Consortium (NNX10AK74H), and a postdoctoral fellowship (RLZ) and graduate student fellowship (AJM) from NSF-funded Center for Dark Energy Biosphere investigations. This is CDEBI contribution # XXX.

We thank Frances O. Mark for valuable technical assistance.

## Author contributions

J.B, I.F and R.L.Z. conceived the study. R.L.Z recovered the genomes from the metagenomic datasets. I.F, R.L.Z analyzed the genomic data. A.M and C.H collected the samples. R.Z revised the thermodynamic calculations and I.F., J.B and R.L.Z. wrote the manuscript with input from all authors. All documents were edited and approved by all authors.

## Competing interests

The other authors declare no competing interests.

## Data availability

The genomes of this study have been made publicly available on GenBank under BioSample accessions (SAMN12695919-SAMN12695949). Metagenomes can be located at NCBI GenBank SRA under project PRJEB11766.

## References

[1] A. Teske and K. B. Sørensen, “Uncultured archaea in deep marine subsurface sediments: have we caught them all?,” ISME J., vol. 2, no. 1, pp. 3–18, Jan. 2008.

[2] J. F. Biddle et al., “Heterotrophic Archaea dominate sedimentary subsurface ecosystems off Peru,” Proc. Natl. Acad. Sci. U. S. A., vol. 103, no. 10, pp. 3846–3851, Mar. 2006.

[3] A. Schippers et al., “Prokaryotic cells of the deep sub-seafloor biosphere identified as living bacteria,” Nature, vol. 433, no. 7028, pp. 861–864, Feb. 2005.

[4] J. S. Lipp, Y. Morono, F. Inagaki, and K.-U. Hinrichs, “Significant contribution of Archaea to extant biomass in marine subsurface sediments,” Nature, vol. 454, no. 7207, pp. 991–994, Aug. 2008.

[5] J. Buongiorno et al., “Interlaboratory quantification of Bacteria and Archaea in deeply buried sediments of the Baltic Sea (IODP Expedition 347),” FEMS Microbiol. Ecol., vol. 93, no. 3, 01 2017.

[6] J. F. Biddle, S. Fitz-Gibbon, S. C. Schuster, J. E. Brenchley, and C. H. House, “Metagenomic signatures of the Peru Margin subseafloor biosphere show a genetically distinct environment,” Proc. Natl. Acad. Sci. U. S. A., vol. 105, no. 30, pp. 10583–10588, Jul. 2008.

[7] N. H. Youssef, C. Rinke, R. Stepanauskas, I. Farag, T. Woyke, and M. S. Elshahed, “Insights into the metabolism, lifestyle and putative evolutionary history of the novel archaeal phylum ‘Diapherotrites,’” ISME J., vol. 9, no. 2, pp. 447–460, Feb. 2015.

[8] I. F. Farag, N. H. Youssef, and M. S. Elshahed, “Global Distribution Patterns and Pangenomic Diversity of the Candidate Phylum ‘Latescibacteria’ (WS3),” Appl. Environ. Microbiol., vol. 83, no. 10, 15 2017.

[9] K. Kubo, K. G. Lloyd, J. F Biddle, R. Amann, A. Teske, and K. Knittel, “Archaea of the Miscellaneous Crenarchaeotal Group are abundant, diverse and widespread in marine sediments,” ISME J., vol. 6, no. 10, pp. 1949–1965, Oct. 2012.

[10] C. Rinke et al., “Insights into the phylogeny and coding potential of microbial dark matter,” Nature, vol. 499, no. 7459, pp. 431–437, Jul. 2013.

[11] A. Spang et al., “Complex archaea that bridge the gap between prokaryotes and eukaryotes,” Nature, vol. 521, no. 7551, pp. 173–179, May 2015.

[12] L. A. Hug et al., “A new view of the tree of life,” Nat. Microbiol., vol. 1, p. 16048, Apr. 2016.

[13] C. J. Castelle and J. F. Banfield, “Major New Microbial Groups Expand Diversity and Alter our Understanding of the Tree of Life,” Cell, vol. 172, no. 6, pp. 1181–1197, 08 2018.

[14] K. Anantharaman et al., “Thousands of microbial genomes shed light on interconnected biogeochemical processes in an aquifer system,” Nat. Commun., vol. 7, p. 13219, 24 2016.

[15] K. Zaremba-Niedzwiedzka et al., “Asgard archaea illuminate the origin of eukaryotic cellular complexity,” Nature, vol. 541, no. 7637, pp. 353–358, 19 2017.

[16] A. Spang et al., “Proposal of the reverse flow model for the origin of the eukaryotic cell based on comparative analyses of Asgard archaeal metabolism,” Nat. Microbiol., vol. 4, no. 7, pp. 1138–1148, 2019.

[17] F. MacLeod, G. S. Kindler, H. L. Wong, R. Chen, and B. P. Burns, “Asgard archaea: Diversity, function, and evolutionary implications in a range of microbiomes,” AIMS Microbiol., vol. 5, no. 1, pp. 48–61, 2019.

[18] K. W. Seitz et al., “Asgard archaea capable of anaerobic hydrocarbon cycling,” Nat. Commun., vol. 10, no. 1, p. 1822, 23 2019.

[19] Z. Zhou, J. Pan, F. Wang, J.-D. Gu, and M. Li, “Bathyarchaeota: globally distributed metabolic generalists in anoxic environments,” FEMS Microbiol. Rev., vol. 42, no. 5, pp. 639–655, 01 2018.

[20] P. N. Evans et al., “Methane metabolism in the archaeal phylum Bathyarchaeota revealed by genome-centric metagenomics,” Science, vol. 350, no. 6259, pp. 434–438, Oct. 2015.

[21] Y. He et al., “Genomic and enzymatic evidence for acetogenesis among multiple lineages of the archaeal phylum Bathyarchaeota widespread in marine sediments,” Nat. Microbiol., vol. 1, no. 6, p. 16035, 04 2016.

[22] Amanda Martino, Matthew E. Rhodes, Rosa León-Zayas, Isabella E. Valente, Jennifer F. Biddle and Christopher H. House, “Microbial Diversity in Sub-Seafloor Sediments from the Costa Rica Margin,” Geosciences, vol. 9, p. 218, 2019.

[23] S. Podell and T. Gaasterland, “DarkHorse: a method for genome-wide prediction of horizontal gene transfer,” Genome Biol., vol. 8, no. 2, p. R16, 2007.

[24] L. A. Levin et al., “A hydrothermal seep on the Costa Rica margin: middle ground in a continuum of reducing ecosystems,” Proc. Biol. Sci., vol. 279, no. 1738, pp. 2580–2588, Jul. 2012.

[25] R. U. Meckenstock et al., “Anaerobic Degradation of Benzene and Polycyclic Aromatic Hydrocarbons,” J. Mol. Microbiol. Biotechnol., vol. 26, no. 1-3, pp. 92–118, 2016.

[26] M. H. Stagars, S. E. Ruff, R. Amann, and K. Knittel, “High Diversity of Anaerobic Alkane-Degrading Microbial Communities in Marine Seep Sediments Based on (1-methylalkyl)succinate Synthase Genes,” Front. Microbiol., vol. 6, p. 1511, 2015.

[27] B. Song and B. B. Ward, “Genetic diversity of benzoyl coenzyme A reductase genes detected in denitrifying isolates and estuarine sediment communities,” Appl. Environ. Microbiol., vol. 71, no. 4, pp. 2036–2045, Apr. 2005.

[28] M. J. López Barragán et al., “The bzd gene cluster, coding for anaerobic benzoate catabolism, in Azoarcus sp. strain CIB,” J. Bacteriol., vol. 186, no. 17, pp. 5762–5774, Sep. 2004.

[29] J.-M. Huang, B. J. Baker, J.-T. Li, and Y. Wang, “New Microbial Lineages Capable of Carbon Fixation and Nutrient Cycling in Deep-Sea Sediments of the Northern South China Sea,” Appl. Environ. Microbiol., vol. 85, no. 15, Aug. 2019.

[30] K. Mendler, H. Chen, D. H. Parks, B. Lobb, L. A. Hug, and A. C. Doxey, “AnnoTree: visualization and exploration of a functionally annotated microbial tree of life,” Nucleic Acids Res., vol. 47, no. 9, pp. 4442–4448, May 2019.

[31] C. Greening et al., “Genomic and metagenomic surveys of hydrogenase distribution indicate H2 is a widely utilised energy source for microbial growth and survival,” ISME J., vol. 10, no. 3, pp. 761–777, Mar. 2016.

[32] A. Kouzuma, S. Kato, and K. Watanabe, “Microbial interspecies interactions: recent findings in syntrophic consortia,” Front. Microbiol., vol. 6, p. 477, 2015.

[33] S. Pirbadian et al., “Shewanella oneidensis MR-1 nanowires are outer membrane and periplasmic extensions of the extracellular electron transport components,” Proc. Natl. Acad. Sci. U. S. A., vol. 111, no. 35, pp. 12883–12888, Sep. 2014.

[34] S. L. McCrindle, U. Kappler, and A. G. McEwan, “Microbial dimethylsulfoxide and trimethylamine-N-oxide respiration,” Adv. Microb. Physiol., vol. 50, pp. 147–198, 2005.

[35] P. N. Evans et al., “An evolving view of methane metabolism in the Archaea,” Nat. Rev. Microbiol., vol. 17, no. 4, pp. 219–232, Apr. 2019.

[36] P. S. Adam, G. Borrel, and S. Gribaldo, “Evolutionary history of carbon monoxide dehydrogenase/acetyl-CoA synthase, one of the oldest enzymatic complexes,” Proc. Natl. Acad. Sci. U. S. A., vol. 115, no. 6, pp. E1166–E1173, 06 2018.

[37] L. A. Hug et al., “Community genomic analyses constrain the distribution of metabolic traits across the Chloroflexi phylum and indicate roles in sediment carbon cycling,” Microbiome, vol. 1, no. 1, p. 22, Aug. 2013.

[38] M. Mehrshad et al., “Hidden in plain sight-highly abundant and diverse planktonic freshwater Chloroflexi,” Microbiome, vol. 6, no. 1, p. 176, 02 2018.

[39] B. B. Jørgensen and A. Boetius, “Feast and famine--microbial life in the deep-sea bed,” Nat. Rev. Microbiol., vol. 5, no. 10, pp. 770–781, Oct. 2007.

[40] X. Dong et al., “Metabolic potential of uncultured bacteria and archaea associated with petroleum seepage in deep-sea sediments,” Nat. Commun., vol. 10, no. 1, p. 1816, 18 2019.

[41] Y.-F. Liu et al., “Anaerobic hydrocarbon degradation in candidate phylum ‘Atribacteria’ (JS1) inferred from genomics,” ISME J., Jun. 2019.

[42] M. A. Lever, “Acetogenesis in the energy-starved deep biosphere - a paradox?,” Front. Microbiol., vol. 2, p. 284, 2011.

[43] M. A. Mausz and Y. Chen, “Microbiology and Ecology of Methylated Amine Metabolism in Marine Ecosystems,” Curr. Issues Mol. Biol., vol. 33, pp. 133–148, Jun. 2019.

[44] Vannucchi, P.; Ujiie, K.; Stroncik, N.; IODP Exp. 334 Scientific Party; Yatheesh, V., “IODP expedition 334: An investigation of the sedimentary record, fluid flow and state of stress on top of the seismogenic zone of an erosive subduction margin,” Sci. Drill., vol. vol.15, pp. 23–30, 2013.

[45] Y. Peng, H. C. M. Leung, S. M. Yiu, and F. Y. L. Chin, “IDBA-UD: a de novo assembler for single-cell and metagenomic sequencing data with highly uneven depth,” Bioinforma. Oxf. Engl., vol. 28, no. 11, pp. 1420–1428, Jun. 2012.

[46] Y.-W. Wu, Y.-H. Tang, S. G. Tringe, B. A. Simmons, and S. W. Singer, “MaxBin: an automated binning method to recover individual genomes from metagenomes using an expectation-maximization algorithm,” Microbiome, vol. 2, p. 26, 2014.

[47] C. C. Laczny et al., “VizBin - an application for reference-independent visualization and human-augmented binning of metagenomic data,”Microbiome, vol. 3, no. 1, p. 1, 2015.

[48] D. H. Parks, M. Imelfort, C. T. Skennerton, P. Hugenholtz, and G. W. Tyson, “CheckM: assessing the quality of microbial genomes recovered from isolates, single cells, and metagenomes,” Genome Res., vol. 25, no. 7, pp. 1043–1055, Jul. 2015.

[49] M. R. Olm, C. T. Brown, B. Brooks, and J. F. Banfield, “dRep: a tool for fast and accurate genomic comparisons that enables improved genome recovery from metagenomes through de-replication,” ISME J., vol. 11, no. 12, pp. 2864–2868, 2017.

[50] A. E. Darling, G. Jospin, E. Lowe, F. A. Matsen, H. M. Bik, and J. A. Eisen, “PhyloSift: phylogenetic analysis of genomes and metagenomes,” PeerJ, vol. 2, p. e243, 2014.

[51] T. Seemann, “Prokka: rapid prokaryotic genome annotation,” Bioinforma. Oxf. Engl., vol. 30, no. 14, pp. 2068–2069, Jul. 2014.

[52] D. Hyatt, G.-L. Chen, P. F. Locascio, M. L. Land, F. W. Larimer, and L. J. Hauser, “Prodigal: prokaryotic gene recognition and translation initiation site identification,” BMC Bioinformatics, vol. 11, p. 119, Mar. 2010.

[53] L.-T. Nguyen, H. A. Schmidt, A. von Haeseler and B. Q. Minh, “IQ-TREE: a fast and effective stochastic algorithm for estimating maximum-likelihood phylogenies,” Mol. Biol. Evol., vol. 32, no. 1, pp. 268–274, Jan. 2015.

[54] M. A. Miller, W. Pfeiffer, and T. Schwartz, “Creating the CIPRES Science Gateway for inference of large phylogenetic trees,” 2010 Gatew. Comput. Environ. Workshop GCE, pp. 1–8, 2010.

[55] D. T. Hoang, O. Chernomor, A. von Haeseler, B. Q. Minh, and L. S. Vinh, “UFBoot2: Improving the Ultrafast Bootstrap Approximation,” Mol. Biol. Evol., vol. 35, no. 2, pp. 518–522, 01 2018.

[56] L. S. Johnson, S. R. Eddy, and E. Portugaly, “Hidden Markov model speed heuristic and iterative HMM search procedure,” BMC Bioinformatics, vol. 11, p. 431, Aug. 2010.

[57] M. Kanehisa, Y. Sato, and K. Morishima, “BlastKOALA and GhostKOALA: KEGG Tools for Functional Characterization of Genome and Metagenome Sequences,” J. Mol. Biol., vol. 428, no. 4, pp. 726–731, Feb. 2016.

[58] Y. Yin, X. Mao, J. Yang, X. Chen, F. Mao, and Y. Xu, “dbCAN: a web resource for automated carbohydrate-active enzyme annotation,” Nucleic Acids Res., vol. 40, no. Web Server issue, pp. W445–451, Jul. 2012.

[59] J. J. Almagro Armenteros et al., “SignalP 5.0 improves signal peptide predictions using deep neural networks,” Nat. Biotechnol., vol. 37, no. 4, pp. 420–423, 2019.

[60] R. C. Edgar, “Search and clustering orders of magnitude faster than BLAST,” Bioinforma. Oxf. Engl., vol. 26, no. 19, pp. 2460–2461, Oct. 2010.

[61] N. D. Rawlings, A. J. Barrett, P. D. Thomas, X. Huang, A. Bateman, and R. D. Finn, “The MEROPS database of proteolytic enzymes, their substrates and inhibitors in 2017 and a comparison with peptidases in the PANTHER database,” Nucleic Acids Res., vol. 46, no. D1, pp. D624–D632, 04 2018.

[62] M. H. Saier, V. S. Reddy, B. V. Tsu, M. S. Ahmed, C. Li, and G. Moreno-Hagelsieb, “The Transporter Classification Database (TCDB): recent advances,” Nucleic Acids Res., vol. 44, no. D1, pp. D372–379, Jan. 2016.

[63] P. Jones et al., “InterProScan 5: genome-scale protein function classification,” Bioinforma. Oxf. Engl., vol. 30, no. 9, pp. 1236–1240, May 2014.

[64] R. C. Edgar, “MUSCLE: multiple sequence alignment with high accuracy and high throughput,” Nucleic Acids Res., vol. 32, no. 5, pp. 1792–1797, 2004.

[65] A. Flamholz, E. Noor, A. Bar-Even, and R. Milo, “eQuilibrator--the biochemical thermodynamics calculator,” Nucleic Acids Res., vol. 40, no. Database issue, pp. D770–775, Jan. 2012.

[66] D. E. LaRowe and J. P. Amend, “Catabolic rates, population sizes and doubling/replacement times of microorganisms in natural settings,” Am. J. Sci., vol. 315, pp. 167–203, Mar. 2015.

[67] J. M. Dick, “Calculation of the relative metastabilities of proteins using the CHNOSZ software package,” Geochem. Trans., vol. 9, p. 10, Oct. 2008.

[68] H. C. Helgeson, “Thermodynamics of hydrothermal systems at elevated temperatures and pressures,” Am. J. Sci., vol. 267, pp. 729–804, Sep. 1969.

