## Supplementary figures (1-7), supplementary tables (1-2) for "Genomic expansion of archaeal lineages resolved from deep Costa Rica sediments"

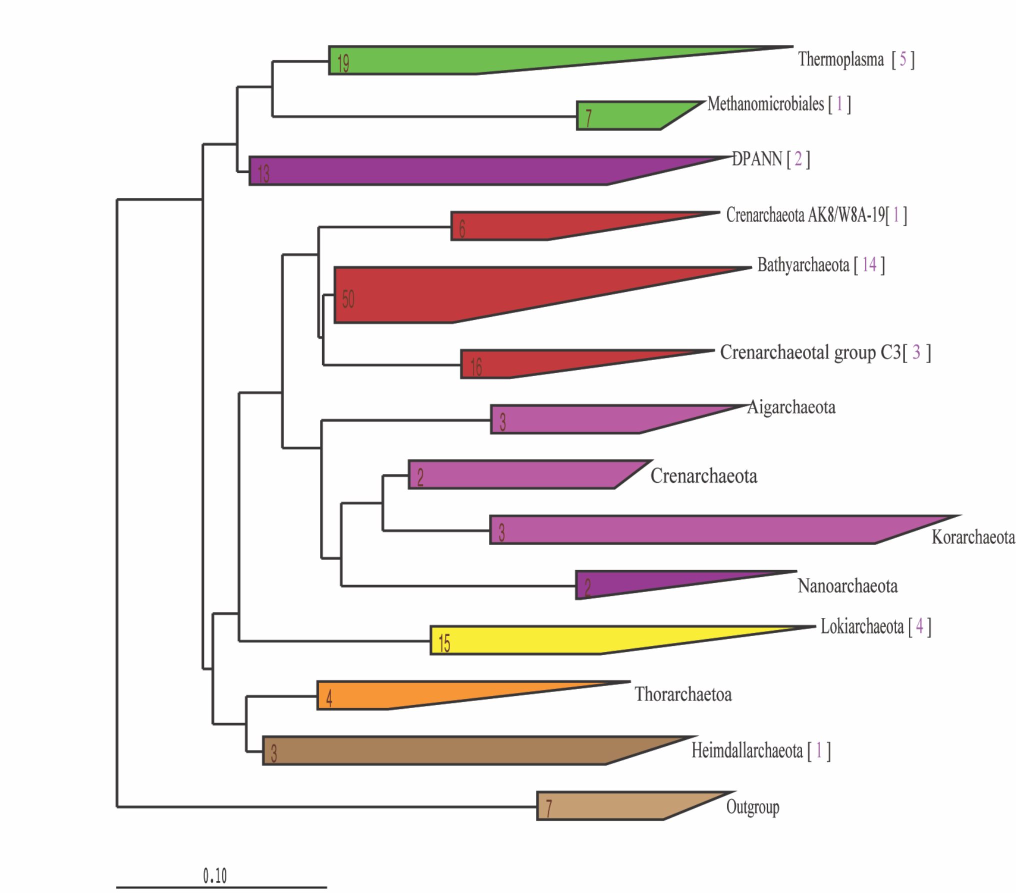


**Figure S1. Phylogenetic placement of the 16S rRNA gene sequences recovered from Costa Rica metagenomes.** The maximum likelihood phylogenetic tree was calculated using ARB. Numbers in the parentheses are the number of 16S rRNA gene sequences recovered from CR metagenomes included in each phylogenetic group.

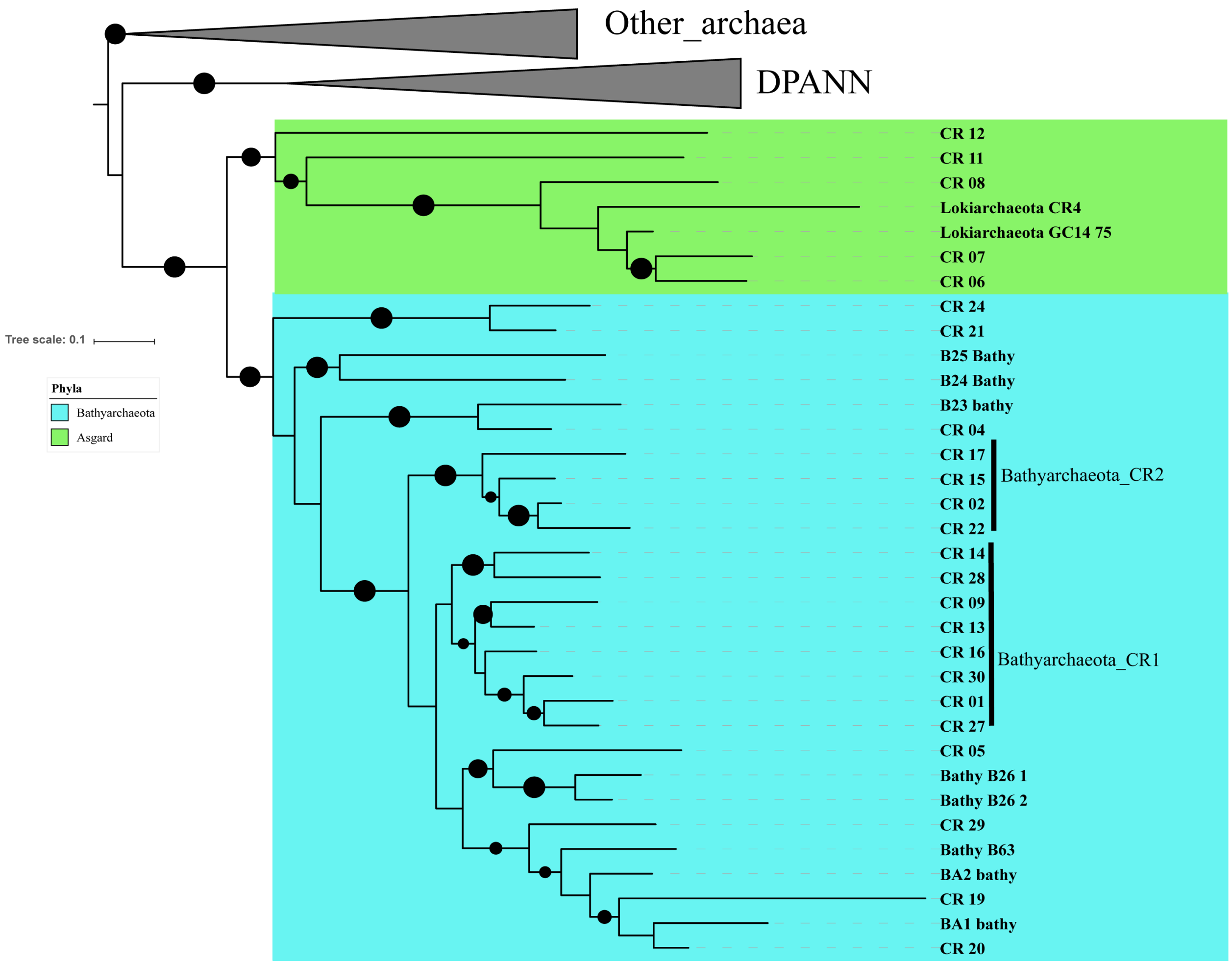


**Figure S2. (A) Phylogenetic placement of the Costa Rica *Bathyarchaeota* and *Lokiarchaeota* MAGs.** The maximum-likelihood phylogenetic tree was calculated based on the concatenation of 16 ribosomal proteins (L2, L3, L4, L5, L6, L14, L15, L16, L18, L22, L24, S3, S8, S10, S17, and S19). “Bathyarchaeota_CR1” and “Bathyarchaeota_CR2” represent two novel classes within the phylum *Bathyarchaeota.* The relationships were inferred using the best fit substitution model (WAG+F+R6) and nodes with bootstrap support >80% were marked by black circles. Scale bar indicates substitutions per site.

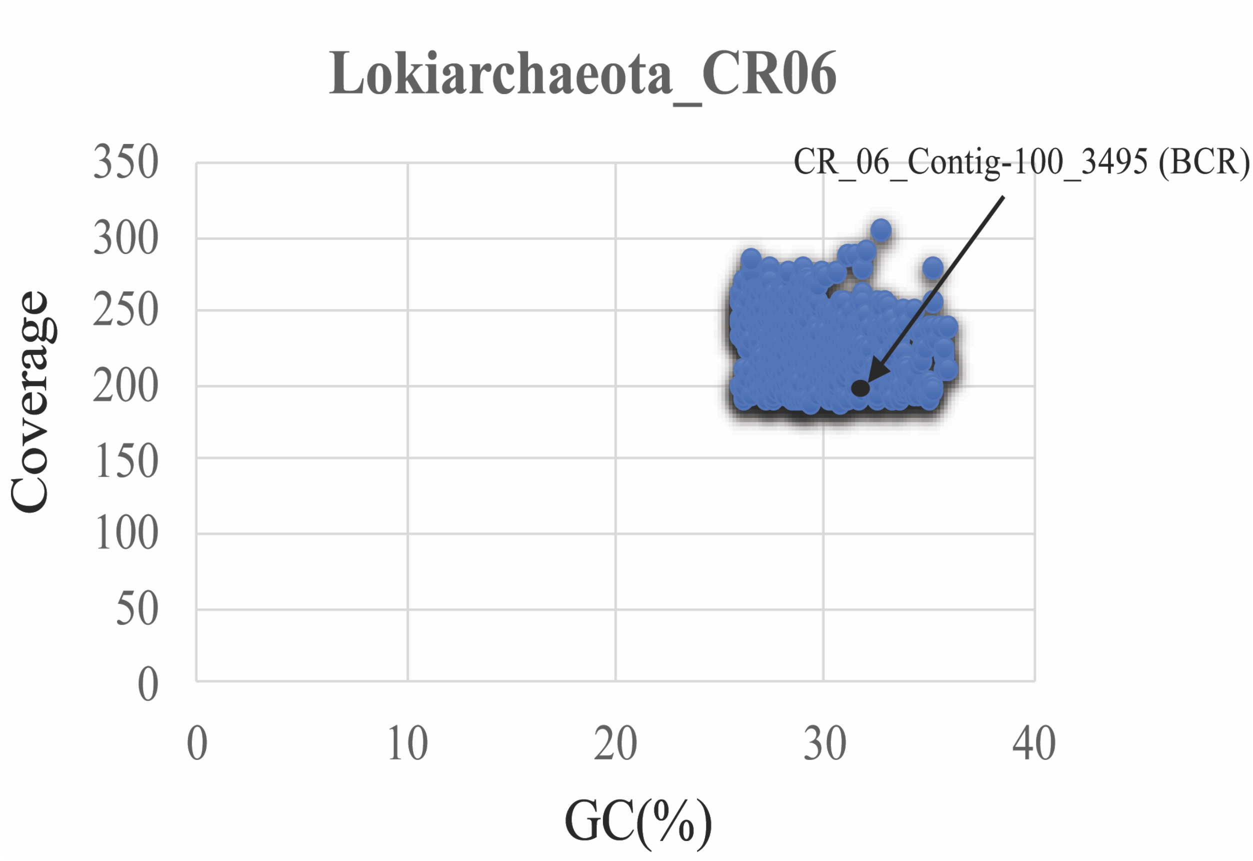


**Figure S3. Coverage and GC (%) range of the contigs within Lokiarchaeota (CR_06) MAG.** CR_06_Contig-100_3495 encodes for the BCR operon and is highlighted by the black dot.

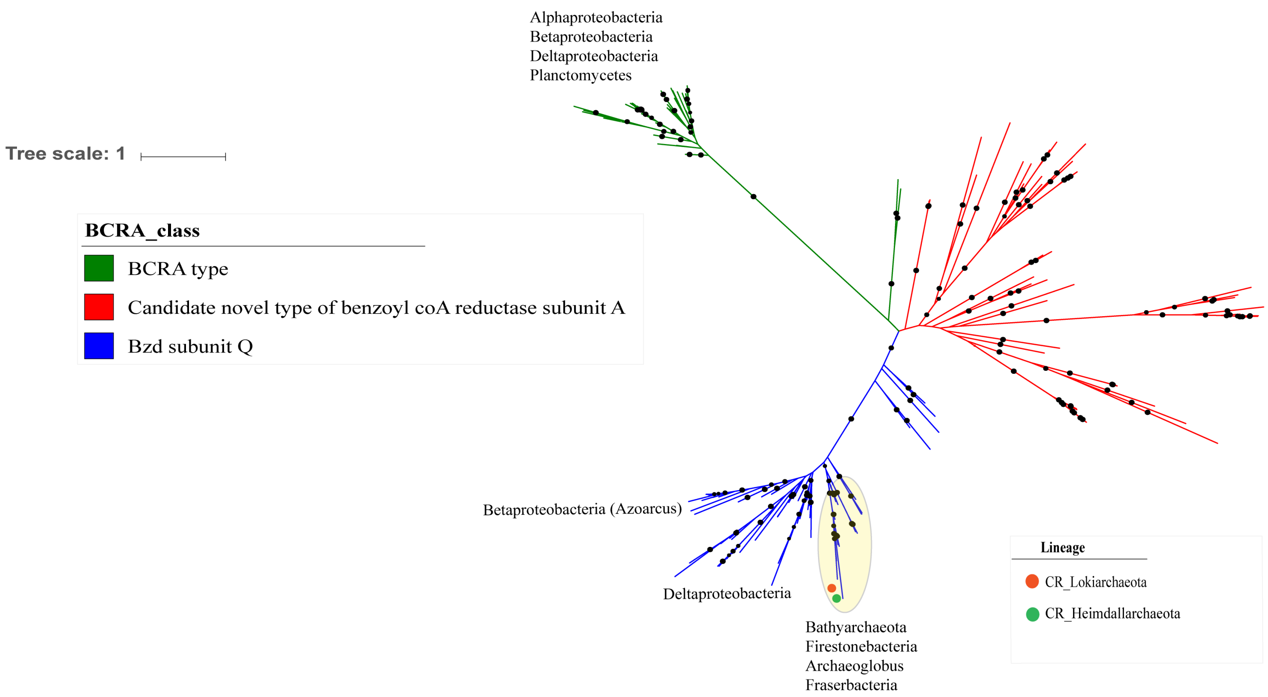


**Figure S4. Maximum likelihood tree of the benzoyl-CoA reductase subunit A.** The tree was calculated using the best fit substitution model (LG+R6) that describes the evolutionary relationships between BCRA families implemented in IQ-Tree [1]. The tree was made using reference sequences under the KEGG entry (K04114) collected from AnnoTree [2] and branch location was tested using 1000 ultrafast bootstraps and approximate Bayesian computation, branches with bootstrap support >80% were marked by black circles. Blue and green clades highlight sequences belong to Bzd_Q and BCR_A subfamilies, respectively. Scale bar indicates substitutions per site. Sequences from CR_*Lokiarchaeota* MAGs were marked with red circles, and CR_*Heimdallarchaeota* was marked with green. Candidate novel clades present in the Costa Rica metagenomic datasets were colored red.

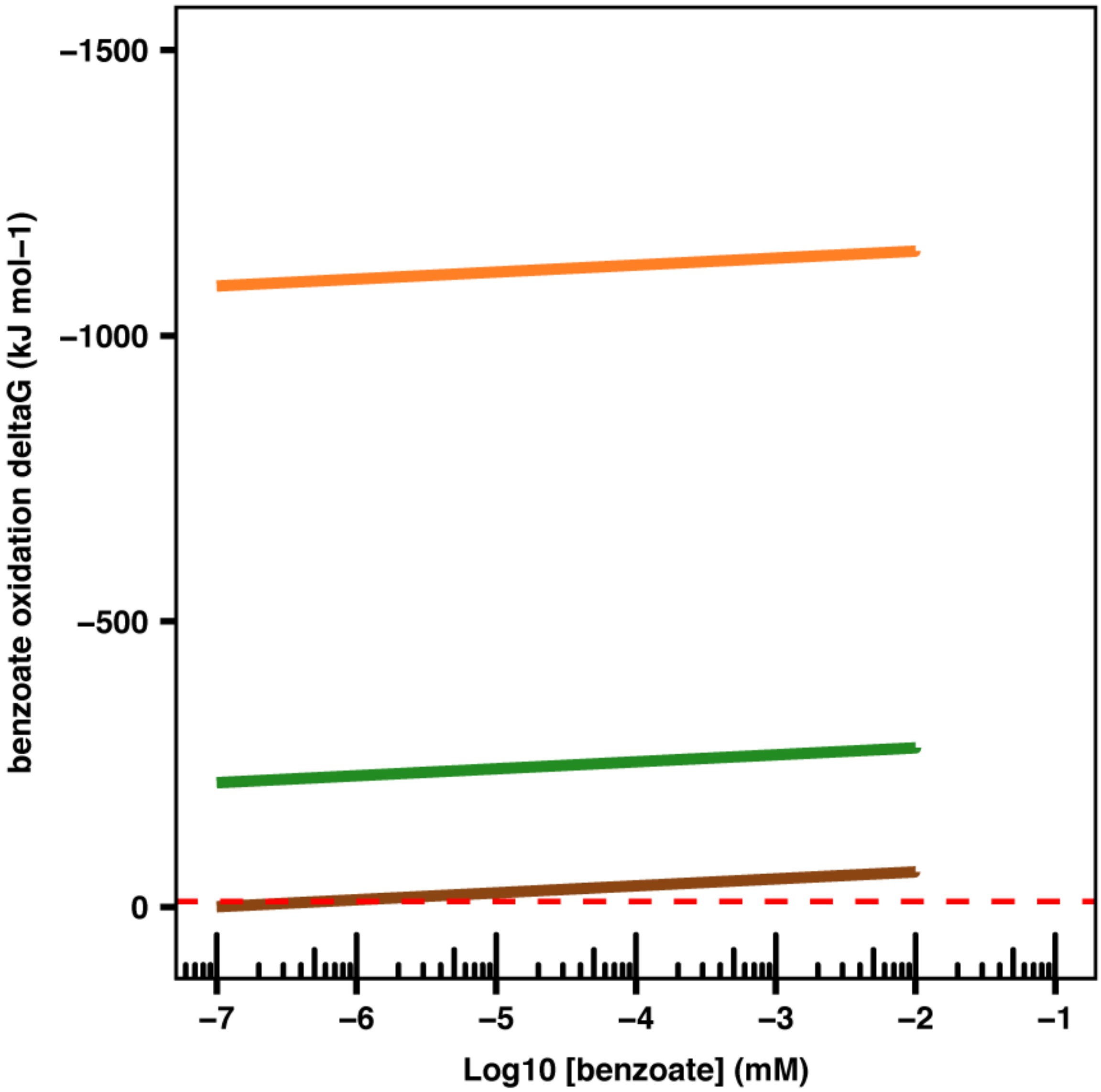


**Figure S5. Gibbs free energy of the proposed catabolic reactions of Lokiarchaeota (CR_06) under the near in situ condition.** Gibbs free energy of benzoate oxidation coupled to the reduction of nitrate (brown line), nitrite (orange line), and sulfite (green line), with the benzoate concentration varying in a wide range ($10^{-4}$–10 µM). Benzoate oxidation coupled to sulfate reduction was calculated to be thermodynamic unfavorable, both under standard condition and the in-situ condition in subseafloor sediments at Costa Rica Margin. The red dashed line indicate the theoretical minimal energy quantum of life [-10 kJ $mol^{-1}$][3]

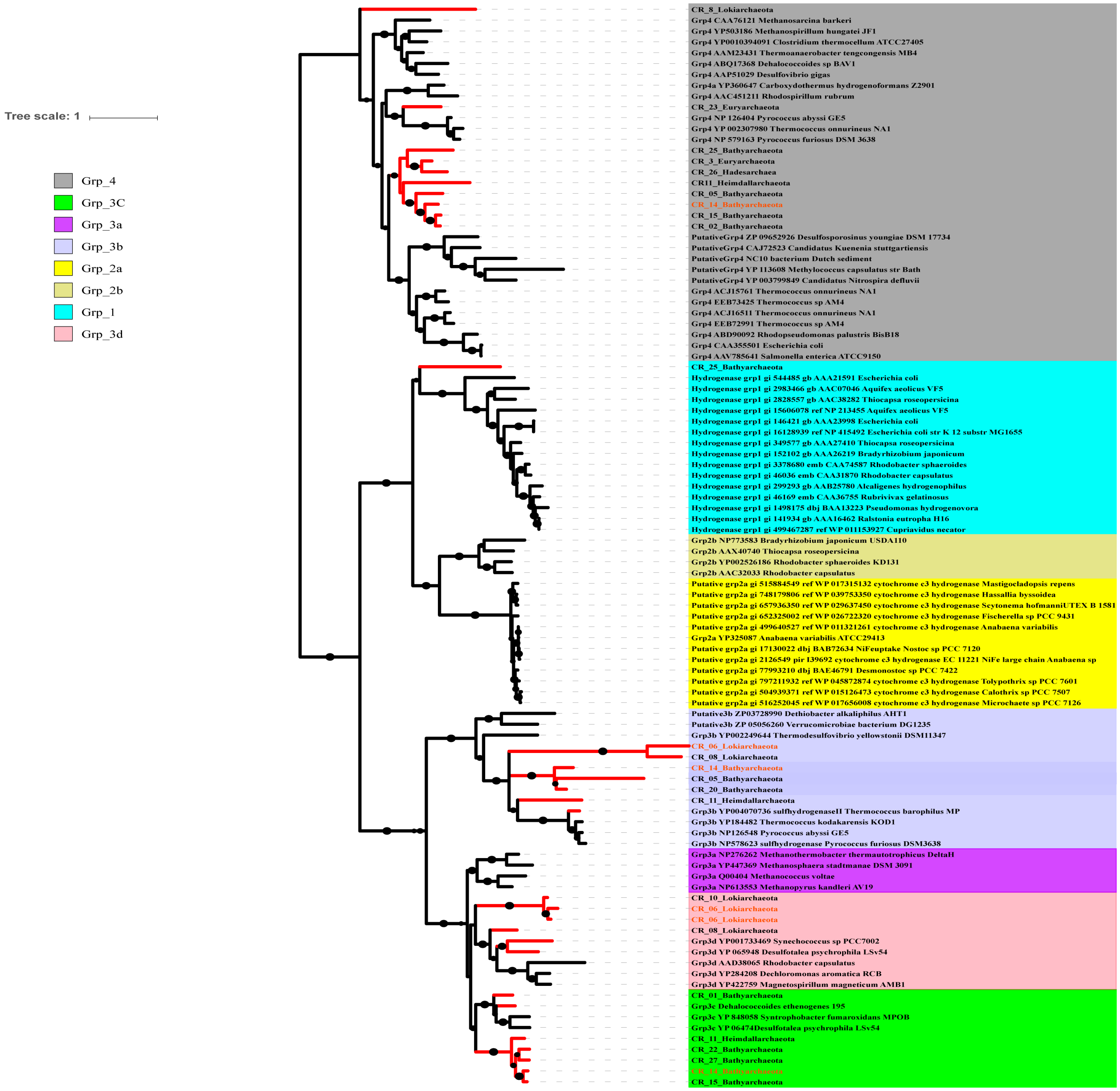


**Figure S6. Maximum likelihood phylogenetic tree of the [NiFe] hydrogenases recovered from CR_MAGs.** The relationships were inferred using the best fit substitution model (LG+R6) and nodes with bootstrap support >80% were marked by black circles. Scale bar indicates substitutions per site. Hydrogenase protein sequences from CR_MAGS are highlighted with red branches. Reference sequences used were collected from (Greening et al. 2016)[4].

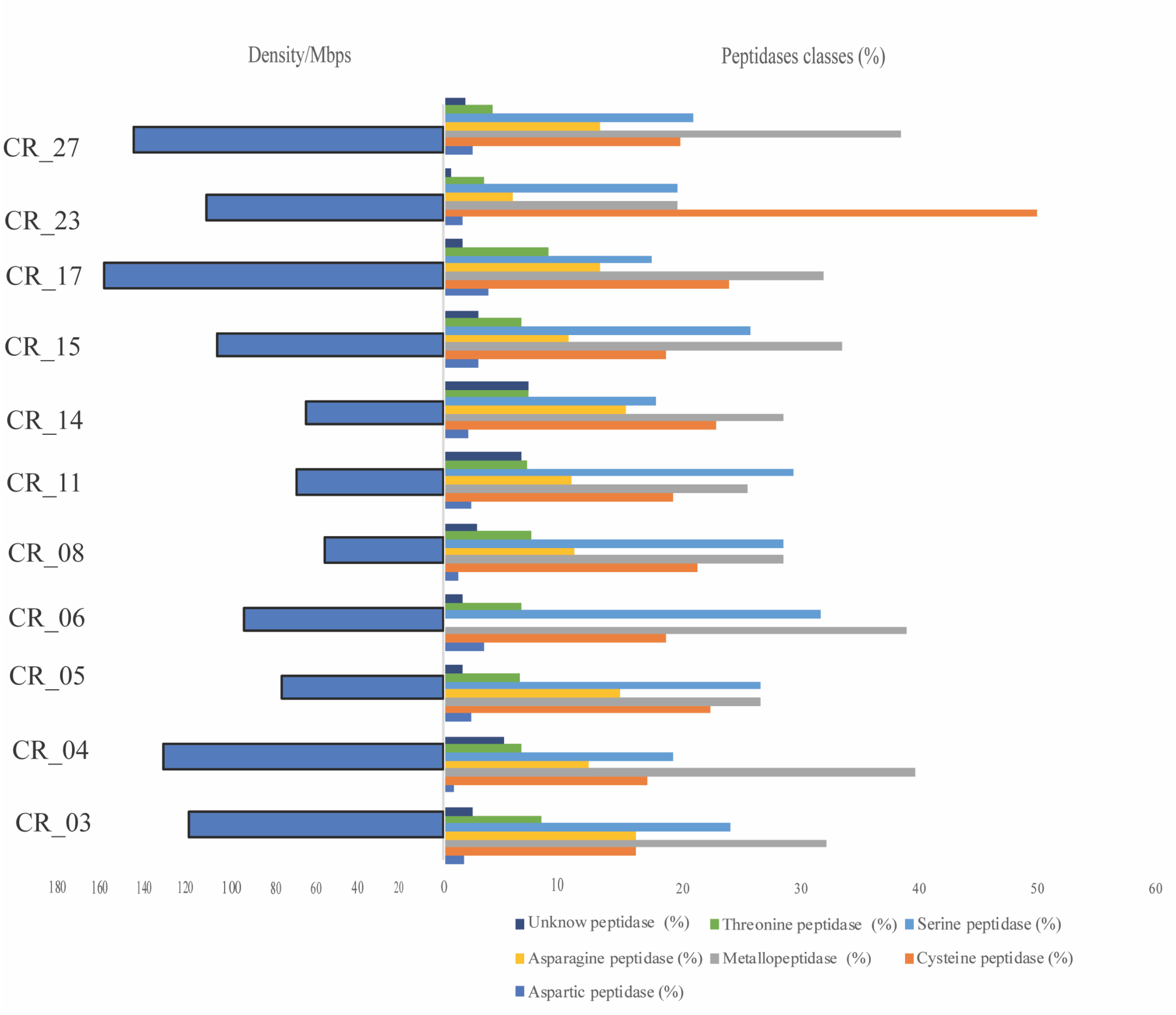


**Figure S7.** (Left panel) Relative densities (numbers per 1 Mb) of peptidases in the CR_MAGs with >60% completeness. (Right panel) abundance percentages of each peptidase class within different Costa Rica MAGs.

**Table S1. Gibbs free energy of couple redox reactions proposed for the CR_*Lokiarchaeota* MAG (CR_06).**

| Reaction | Equation | ΔG° (kJ $mol^{-1}$) |
|---|---|---|
| **Benzoate oxidation coupled to nitrate reduction to nitrite** | $[C_7H_5O^{2-}] + 3[NO^{3-}] + 9[H_2O]$ -> $3[NO_2^-] + 7[CO_2] + 24\ e^-$ | -119.9 |
| **Benzoate oxidation coupled to nitrite reduction to ammonia** | $[C_7H_5O^{2-}] + 3[NO_2^-] + 6[H_2O]$ -> $3[NH_4^+] + 7[CO2] + 12\ e^-$ | -1206.3 |
| **Benzoate oxidation coupled to sulfate reduction** | $[C_7H_5O^{2-}] + 3[SO_4^{-2}] + 9[H_2O]$ -> $3[SO_3^{-2}] + 7[CO_2] + 24\ e^-$ | 416.1 |
| **Benzoate oxidation coupled to sulfite reduction to hydrogen sulfide** | $[C_7H_5O^{2-}] + 3[SO_3^{-2}] + 3[H2O]$ -> $3[H_2S] + 7[CO2] + 12\ e^-$ | -373.6 |
| **Benzoate oxidation to acetate** | $2[C_7H_5O^{2-}] + 10[H2O]$ -> $7[C_2H_3O^{2-}] + 2\ e^-$ | 196.3 |

**Table S2. ANI Comparisons between Costa Rica *Bathyarchaeota* MAG CR_14 against reference *Bathyarchaeota* MAGs.**

| CR genome | Reference genome | Distance | Similarity |
|---|---|---|---|
| **CR_14** | B23 | 0.295981 | 0.704019 |
| **CR_14** | B25 | 0.295981 | 0.704019 |
| **CR_14** | B63 | 1 | 0 |
| **CR_14** | BA2 | 0.295981 | 0.704019 |
| **CR_14** | B24 | 0.295981 | 0.704019 |
| **CR_14** | B26 | 1 | 0 |
| **CR_14** | BA1 | 1 | 0 |
| **CR_14** | B26 | 1 | 0 |

**Table S3. ANI comparisons between CR MAGs**

**Table S4. Function annotation of CR_06 proteins using KEGG database**

**Table S5. Function annotation of CR_14 proteins using KEGG database**

**Table S6. HMM searches against key metabolic genes to validate the presence/absence of the pathways in CR-metagenomes**

**Tables S3, S4, S5 and S6 are provided as a separate excel spreadsheet.**

### References


[1] L.-T. Nguyen, H. A. Schmidt, A. von Haeseler, and B. Q. Minh, "IQ-TREE: a fast and effective stochastic algorithm for estimating maximum-likelihood phylogenies," *Mol. Biol. Evol.*, vol. 32, no. 1, pp. 268–274, Jan. 2015.

[2] K. Mendler, H. Chen, D. H. Parks, B. Lobb, L. A. Hug, and A. C. Doxey, "AnnoTree: visualization and exploration of a functionally annotated microbial tree of life," *Nucleic Acids Res.*, vol. 47, no. 9, pp. 4442–4448, May 2019.

[3] V. Müller and V. Hess, "The Minimum Biological Energy Quantum," *Front. Microbiol.*, vol. 8, p. 2019, 2017.

[4] C. Greening *et al.*, "Genomic and metagenomic surveys of hydrogenase distribution indicate H2 is a widely utilised energy source for microbial growth and survival," *ISME J.*, vol. 10, no. 3, pp. 761–777, Mar. 2016.
